# Acetate metabolism during xylose fermentation enhances 3-hydroxypropionic acid production in engineered acid-tolerant *Issatchenkia orientalis*

**DOI:** 10.1101/2025.04.17.649204

**Authors:** Deokyeol Jeong, Dahye Lee, Junli Liu, Soo Rin Kim, Yong-Su Jin, Jikai Zhao, Eun Joong Oh

## Abstract

Efficient bioconversion of acetate-rich lignocellulosic biomass into value-added chemicals remains a major challenge due to the toxicity of acetic acid. In this study, we engineered an acid-tolerant *Issatchenkia orientalis* strain (IoDY01H) capable of producing 3-hydroxypropionic acid (3-HP), a key bioplastic precursor, from glucose, xylose, and acetate. Using a Cas9-based genome editing system with a hygromycin B resistance marker, we introduced heterologous genes encoding xylose utilization and β-alanine-based 3-HP biosynthetic pathways into the *I. orientalis* genome. Metabolomic analysis revealed that acetate supplementation redirected metabolic flux toward amino acid and lipid metabolism while reducing TCA cycle intermediates. Acetate enhanced 3-HP production by promoting accumulation of β-alanine, but also revealed β-alanine–pyruvate aminotransferase as a metabolic bottleneck under acidic conditions. Using pretreated hemp stalk hydrolysate as a feedstock, the engineered strain achieved a 3-HP titer of 8.7 g/L via separate hydrolysis and fermentation (SHF), outperforming simultaneous saccharification and fermentation (SSF). These findings demonstrate the feasibility of producing 3-HP from acetate-rich biomass using engineered non-conventional yeast and highlight *I. orientalis* as a promising microbial chassis for industrial bioconversion.

**Graphical abstract:** 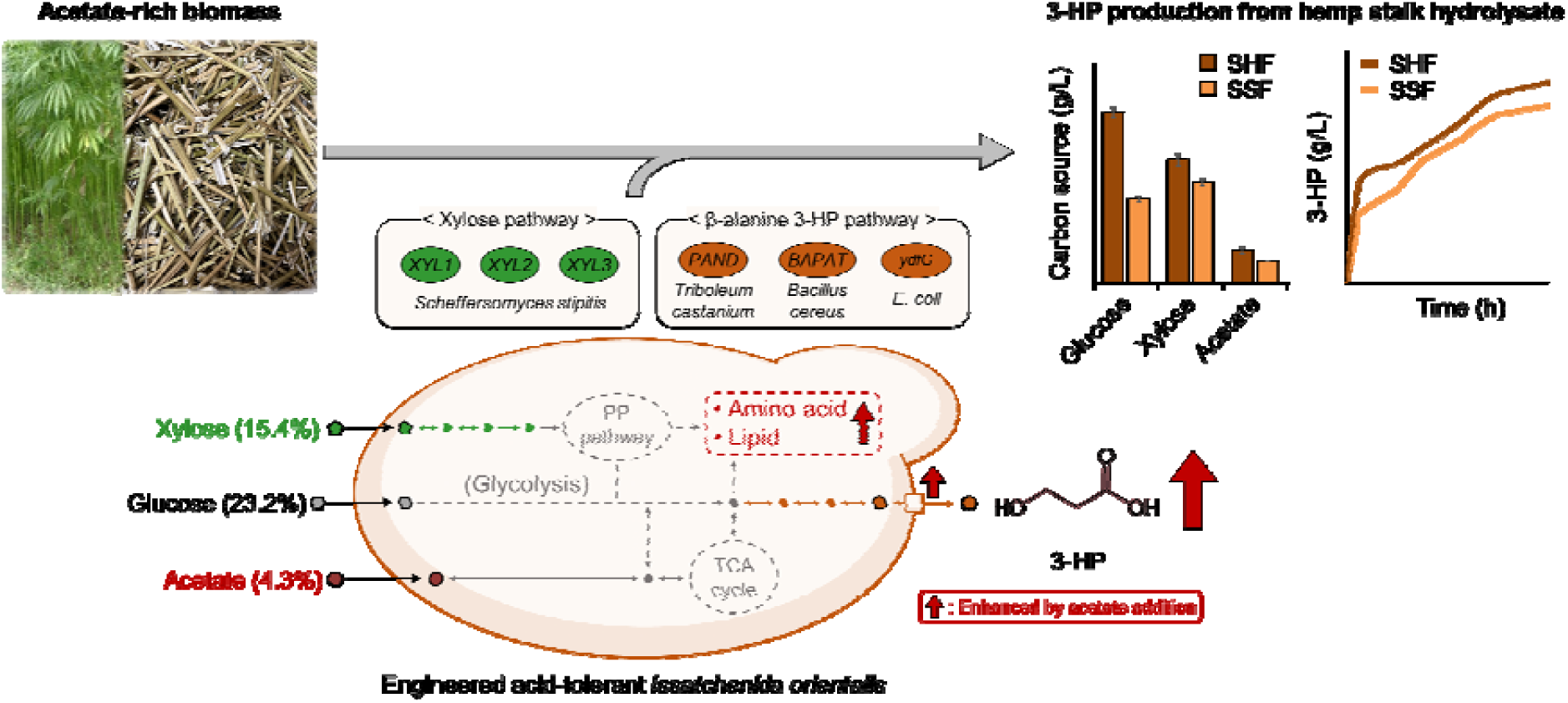

**Highlights:**

- Engineered *I. orientalis* co-utilized xylose and acetate to produce 3-HP.
- Acetate addition enhanced 3-HP production during xylose fermentation.
- Acetate redirected metabolic flux toward 3-HP biosynthesis pathways.
- Engineered *I. orientalis* achieved 8.7 g/L 3-HP from hemp stalk hydrolysate via SHF.

## 1. Introduction

The conversion of renewable biomass into value-added chemicals via microbial fermentation is being developed as a sustainable alternative to conventional petroleum-based production processes to reduce global reliance on fossil fuels and mitigate carbon emissions (Ge et al., 2023; Liang et al., 2024; Volk et al., 2022; Zhou et al., 2024). Lignocellulosic biomass, composed of cellulose, hemicellulose, and lignin, is hydrolyzed to release carbon sources such as glucose, xylose, arabinose, and galactose. Numerous strategies have been developed to efficiently produce value-added products from non-conventional carbon sources such as xylose and arabinose (Feng et al., 2023). However, acetic acid, generated from the acetylation of hemicellulose and lignin in plant cell walls, presents a significant barrier to the widespread utilization of lignocellulosic biomass (Mutyala & Kim, 2022). Rice straw and corn cob residues contain approximately 0.4-3.2% acetic acid, whereas woody biomass sources such as switchgrass, hemp, eucalyptus, and kenaf have higher acetic acid levels ranging from 4.1-6.4% (Lee et al., 2022a). Accumulation of acetic acid inhibits cellular function by inducing intracellular acidification through acetate dissociation and proton (H^+^) release, which increases ATP consumption for proton extrusion, thereby reducing fermentation efficiency (Abbott et al., 2007; Guaragnella & Bettiga, 2021).

Despite advances such as the development of low-acetylation hydrolyzing methods for lignocellulosic biomass, optimization of pretreatment processes, and genetic modification of microorganisms for adaptive evolution and enhanced acetic acid tolerance, acetic acid remains a significant inhibitor in culture media (Mutyala & Kim, 2022; Sun et al., 2021). To address this, strategies for acetate conversion are necessary. Acetate can serve as both an energy and carbon source for the biosynthesis of value-added compounds including ethanol, fatty acids, triacetic acid lactone, vitamin A, lycopene, ethyl acetate, malic acid, mevalonic acid, isobutanol, and succinic acid in model organisms such as *Escherichia coli* and *Saccharomyces cerevisiae* (Gong et al., 2022; Huang et al., 2024; Shi et al., 2021; Sun et al., 2021; Wang et al., 2023). However, the intrinsic acid sensitivity and limited cost-efficiency of these model hosts under low-pH conditions constrain their industrial utility, underscoring the potential of acid-tolerant non-conventional yeasts such as *Issatchenkia orientalis* (Thorwall et al., 2020).

The selection of acid-tolerant microorganisms as industrial workhorse strains offers fermentation advantages under low pH conditions. The multi-tolerant yeast *I. orientalis* can grow at pH 1.5 (Dubinkina et al., 2024) and withstand high temperatures (Seong et al., 2017), showing strong tolerance to furan derivatives and organic acids such as acetic acid (Li et al., 2024; Suthers & Maranas, 2022). Recent strategies leveraging advanced gene editing have facilitated the production of lactic acid (Lee et al., 2024), succinic acid (Tran et al., 2023), itaconic acid (Sun et al., 2020), and citramalate (Wu et al., 2023). The self-buffering capability of *I. orientalis* provides operational cost benefits, including reduced pH adjustment expenses, decreased use of chemicals, lower by-product (e.g., gypsum) accumulation, and minimized wash water requirements (Lee et al., 2024). These advantages contribute to lower global warming potential and reduced fossil energy consumption.

3-hydroxypropionic acid (3-HP) is one of the top 12 platform chemicals that can be produced from biomass via microbial fermentation as identified by the U.S. Department of Energy (Bozell & Petersen, 2010). It serves as a precursor for plastics, resins and other substances (Choi et al., 2015; Choi et al., 2020). Excluding thermodynamically infeasible routes, 3-HP is primarily synthesized via the glycerol, malonyl-CoA, and β-alanine pathways by engineered microorganisms such as *Lactobacillus reuteri*, *E. coli*, *S. cerevisiae*, *Corynebacterium glutamicum*, and *I. orientalis* with production titers reaching up to 125.93 g/L (Barnhart et al., 2017; de Fouchécour et al., 2018; Wang et al., 2024; Zhang et al., 2021). In particular, 3-HP biosynthesis is substrate-dependent, with acetate emerging as a promising carbon source for its production (Kildegaard et al., 2015; Vo & Park, 2024). However, the accumulation of high 3-HP concentrations can cause acidification of the medium, ultimately reducing long-term production efficiency (Qin et al., 2024).

In this study, we engineered an *I. orientalis* strain capable of producing 3-HP from glucose, xylose, and acetate using a hygromycin B antibiotic marker-based Cas9 genetic tool. The strain was designed to detoxify acetate-rich biomass while enabling the utilization of diverse carbon sources. First, we evaluated the strain’s resistance to acetate and 3-HP. Notably, the engineered strain produced 3-HP from xylose, and its production was enhanced by the addition of acetate during xylose fermentation. Intracellular metabolome analysis revealed that the acetate metabolism promoted the accumulation of key intermediates in the 3-HP biosynthetic pathway. Finally, we achieved a 3-HP titer of 8.7 g/L by co-utilizing glucose, xylose, and acetate derived from hemp stalk hydrolysate, a biomass feedstock rich in acetate.

## 2. Materials and methods

### 2.1. Strains and culture conditions

The recombinant yeast strains and plasmids used in this study are listed in Table 1. All yeast strains were cultivated in yeast extract-peptone (YP) medium (10 g/L yeast extract and 20 g/L peptone) supplemented with 20 g/L glucose (YPD) at 30 □ and 250 rpm. *S. cerevisiae* was used to construct expression cassettes for the *I. orientalis* strains. For *S. cerevisiae* transformation, YPD agar supplemented with antibiotics (300 μg/mL G418 sulfate and 300 μg/mL hygromycin B) was used. To assess antibiotic resistance, wild-type *S. cerevisiae* D452-2 and five *I. orientalis* strains (SD108, NRRL Y-27441, ATCC MBY 1358, ATCC 24210, and NRRL YB-373) were cultured on YPD agar containing G418 sulfate or hygromycin B at concentrations of 300, 350, 400, 500, and 600 μg/mL. *E. coli* TOP10 (Invitrogen, Carlsbad, CA, USA) was used for plasmid DNA amplification and cultured in Luria-Bertani medium (5 g/L yeast extract, 10 g/L tryptone, and 10 g/L NaCl) supplemented with 100 μg/mL ampicillin at 37 □ and 250 rpm. All media were sterilized at 121 □ for 15 min, and 20 g/L agar was added when solid medium was required.

**Table 1.** Yeast strains and plasmids used in this study.

| Strains/plasmids | Description/relevant genotype<sup>1)</sup> | Reference |
| --- | --- | --- |
| ***Saccharomyces cerevisiae*** |  |  |
| ScD452-2 | Wild-type; Mata *leu2 his3 ura3* | (Hosaka et al., 1992) |
| D452 *IoTDH3<sub>p</sub>* | ScD452-2 *int#6::IoTDH3<sub>p</sub>-cg#1-IoMDH1<sub>T</sub>* | This study |
| D452 *IoGPM1<sub>p</sub>* | ScD452-2 *int#1::IoGPM1<sub>p</sub>-cg#1-IoPDC1<sub>T</sub>* | This study |
| D452 *IoTEF1<sub>p</sub>* | ScD452-2 *int#1::IoTEF1<sub>p</sub>-cg#1-IoINO1<sub>T</sub>* | This study |
| D452 *IoXYL1* | ScD452-2 *int#6::IoTDH3<sub>p</sub>-XYL1-IoMDH1<sub>T</sub>* | This study |
| D452 *IoXYL2* | ScD452-2 *int#1::IoGPM1<sub>p</sub>-XYL2-IoPDC1<sub>T</sub>* | This study |
| D452 *IoXYL3* | ScD452-2 *int#1::IoTEF1<sub>p</sub>-XYL3-IoINO1<sub>T</sub>* | This study |
| D452 *IoXYL123* | ScD452-2 *int#6::IoTDH3<sub>p</sub>-XYL1-IoMDH1<sub>T</sub>-IoGPM1<sub>p</sub>-XYL2-IoPDC1<sub>T</sub>-IoTEF1<sub>p</sub>-XYL3-IoINO1<sub>T</sub>* | This study |
| D452 *IoTcPAND* | ScD452-2 *int#6::IoTDH3<sub>p</sub>-TcPAND-IoMDH1<sub>T</sub>* | This study |
| D452 *IoBcBAPAT* | ScD452-2 *int#1::IoGPM1<sub>p</sub>-BcBAPAT-IoPDC1<sub>T</sub>* | This study |
| D452 *IoEcydfG* | ScD452-2 *int#1::IoTEF1<sub>p</sub>-EcydfG-IoINO1<sub>T</sub>* | This study |
| D452 *Io3HP* | ScD452-2 *int#6::IoTDH3<sub>p</sub>-TcPAND-IoMDH1<sub>T</sub>-IoGPM1<sub>p</sub>-BcBAPAT-IoPDC1<sub>T</sub>-IoTEF1<sub>p</sub>-EcydfG-IoINO1<sub>T</sub>* | This study |
| ***Issatchenkia orientalis*** |  |  |
| Y-27441 | Wild-type; *Issatchenkia orientalis* NRRL Y-27441 | ARS Culture Collection |
| IoDY01 | Y-27441 *is#1::IoTDH3<sub>p</sub>-XYL1-IoMDH1<sub>T</sub>-IoGPM1<sub>p</sub>-XYL2-IoPDC1<sub>T</sub>-IoTEF1<sub>p</sub>-XYL3-IoINO1<sub>T</sub>* | This study |
| IoDY01H | IoDY01 *is#4::IoTDH3<sub>p</sub>-TcPAND-IoMDH1<sub>T</sub>-IoGPM1<sub>p</sub>-BcBAPAT-IoPDC1<sub>T</sub>-IoTEF1<sub>p</sub>-EcydfG-IoINO1<sub>T</sub>* | This study |
| **Plasmids** |  |  |
| pRS41K-Cas9 | pRS41K, a *natNT* marker, Cas9 | (Kim et al., 2024) |
| pRS42H_cg#1 | pRS42H, a *hph* marker, gRNA block containing created 20-bp sequence for *S. cerevisiae* | (Jeong et al., 2024) |
| pRS42H_INT#1 | pRS42H, a *hph* marker, gRNA block containing for intergenic site#1 of *S. cerevisiae* | (Jeong et al., 2020a) |
| pRS42H_INT#6 | pRS42H, a *hph* marker, gRNA block containing for intergenic site#6 of *S. cerevisiae* | (Jeong et al., 2020a) |
| pCast | pUDP004, a *natNT* marker, iCas9, gRNA block for *I. orientalis* | (Lee et al., 2022b) |
| pCastH | pUDP004, a *hph* marker, iCas9, gRNA block for *I. orientalis* | This study |
| pCastH_IS#1 | pUDP004, a *hph* marker, iCas9, gRNA block containing for intergenic site#1 of *I. orientalis* | This study |
| pCastH_IS#4 | pUDP004, a *hph* marker, iCas9, gRNA block containing for intergenic site#4 of *I. orientalis* | This study |

<sup>1)</sup> The *ydfG* gene derived from *Escherichia coli* TOP10 (*EcydfG*, 3-hydroxypropionate dehydrogenase), and five genes (*XYL1*, xylose reductase; *XYL2*, xylitol dehydrogenase; *XYL3*,
xylulokinase; *TcPAND*, aspartate-1-decarboxylase; *BcBAPAT*, β-alanine-pyruvate aminotransferase) were codon-optimized for *Saccharomyces cerevisiae*.

### 2.2. Construction of expression cassettes for *I. orientalis* in an engineered *S. cerevisiae* strain

All *S. cerevisiae* strains were constructed using Cas9-based genome editing as described previously (Jeong et al., 2020b) and transformed using the LiAc/SS carrier DNA/PEG method (Gietz & Schiestl, 2007). Expression cassettes for the *I. orientalis* strain were assembled in *S. cerevisiae* D452-2 harboring the pRS41K-Cas9 plasmid. Guide RNA (gRNA) plasmids are listed in Table 1, and gRNA sequences are provided in Table S1. The pRS42H_INT#1 plasmid was used to integrate the *IoGPM1*_P_ expression cassette (*IoGPM1*_P_*-cg#1-IoPDC1*_T_) and *IoTEF1*_P_ expression cassette (*IoTEF1*_P_*-cg#1-IoINO1*_T_), while the pRS42H_INT#6 plasmid was used to integrate the *IoTDH3*_P_ expression cassette (*IoTDH3*_P_*-cg#1-IoMDH1*_T_) and additional expression cassettes for the xylose and β-alanine-based 3-HP metabolic pathways. The pRS42H_cg#1 plasmid facilitated the introduction of heterologous genes into the expression cassettes.

Donor DNA fragments were polymerase chain reaction (PCR)-amplified using primers listed in Table S1. For the *IoTDH3*_P_ expression cassette (*IoTDH3*_P_*-cg#1-IoMDH1*_T_), two DNA fragments (*IoTDH3*_P_ and *IoMDH1*_T_) were amplified using the *I. orientalis* NRRL Y-27441 genome and integrated into the intergenic region #6 (*INT#6*) of the *S. cerevisiae* D452-2 strain. Similarly, *IoGPM1*_P_ (*IoGPM1*_P_*-cg#1-IoPDC1*_T_) and *IoTEF1*_P_ (*IoTEF1*_P_*-cg#1-IoINO1*_T_) expression cassettes were integrated into the intergenic region #1 (*INT#1*). For xylose pathway construction, three codon-optimized genes (*XYL1*, *XYL2*, and *XYL3*) were PCR-amplified from the *S. cerevisiae* DY02 genome (Jeong et al., 2020a). The resulting fragments were assembled into the expression cassettes, generating three donor DNAs (*IoTDH3*_P_*-XYL1-IoMDH1*_T_, *IoGPM1*_P_*-XYL2-IoPDC1*_T_, and *IoTEF1*_P_*-XYL3-IoINO1*_T_). These donor DNAs were integrated into the intergenic region #6 (*INT#6*), yielding the D452 *IoXYL123* strain. For 3-HP pathway construction, two codon-optimized *TcPAND* (Accession number PQ863775) and *BcBAPAT* (Accession number PQ863776) genes were synthesized by Twist Biosciences, and the *EcydfG* gene (Accession number NP_416057.1) was PCR-amplified from the *E. coli* TOP10 genome. The three genes (*TcPAND*, *BcBAPAT*, and *EcydfG*) were assembled into donor DNAs (*IoTDH3*_P_*-TcPAND-IoMDH1*_T_, *IoGPM1*_P_*-BcBAPAT-IoPDC1*_T_, and *IoTEF1*_P_*-EcydfG-IoINO1*_T_) and integrated into the intergenic region #6 (*INT#6*) to produce the D452 *Io3HP* strain (Fig. S1B). Correct assembly and genomic integration were validated by colony PCR.

### 2.3. Construction of engineered *I. orientalis* strains

All *I. orientalis* strains were constructed using a Cas9-based genome editing system as described previously (Lee et al., 2022b) with minor modifications. The gRNA plasmids and *I. orientalis* strains, and primers used in this study are listed in Table 1 and Table S1, respectively. To construct the pCastH plasmid, the linearized pCast plasmid lacking the nourseothricin sulfate resistance (*Nat*) marker and hygromycin B (*hyg*) marker fragments were PCR-amplified using OH554/OH555 and OH556/OH557 primers, respectively. The two products were assembled using the HiFi DNA Assembly Cloning Kit (New England Biolabs, E2621L), yielding the pCastH plasmid (Fig. S1A). Subsequently, pCastH plasmids containing the *IS#1* sequence (TAATTTCAACACCTTACTCC) and *IS#4* sequence (AAGGTTTTGCAACTCCCAAG) were constructed as described previously (Lee et al., 2022b), resulting in the pCastH_IS#1 and pCastH_IS#4 plasmids, respectively.

To construct the engineered *I. orientalis* strain capable of xylose consumption, the xylose expression cassette (*IoTDH3_P_-XYL1-IoMDH1_T_-IoGPM1*_P_*-XYL2-IoPDC1*_T_*-IoTEF1*_P_*-XYL3-IoINO1*_T_) was PCR-amplified from the D452 *IoXYL123* strain using primers OH410/OH415. This cassette was then integrated into the intergenic site#1 (*IS#1*) of the *I. orientalis* NRRL Y-27441 strain, resulting in the IoDY01 strain. For the engineered strain producing 3-HP, the 3-HP expression cassette (*IoTDH3*_P_*-TcPAND-IoMDH1*_T_*-IoGPM1*_P_*-BcBAPAT-IoPDC1*_T_*-IoTEF1*_P_*-EcydfG-IoINO1*_T_) was PCR-amplified from the D452 *Io3HP* strain using primers OH774/OH744. This cassette was introduced into the intergenic site #4 (*IS#4*) of the IoDY01 strain, yielding the IoDY01H strain.

All *I. orientalis* strains were transformed by electroporation, with modifications to the protocol previously described (Benatuil et al., 2010; Xi et al., 2021). Briefly, electrocompetent *I. orientalis* cells (∼ 1.6 × 10^9^ cells) were resuspended in 1 mL of electroporation buffer (1 mM CaCl_2_ and 1 M sorbitol). A mixture containing 10 μg of gRNA plasmid and 10 μg donor DNA was added to 200 μL of electroporation buffer, and the volume was adjusted to 400 μL. The reaction was transferred to pre-chilled 2 mm electroporation cuvettes (BTX, BTX620) and incubated on ice for 5 min prior to electroporation. Cells were electroporated at 2.5 kV using an Eppendorf Eporator (Eppendorf, Germany), then immediately transferred to 10 mL of 1:1 (v/v) 1 M sorbitol:YPD recovery medium and incubated at 30 □ with shaking (250 rpm) for 12-18 h. After centrifugation (15,928 × g and 1 min), the supernatant was discarded, and 2 mL of the cell suspension was plated onto YPD agar plates containing 1 M sorbitol and 350 μg/mL hygromycin B. Plates were incubated at 30 □ for 3-4 days. Positive transformants were identified by yeast colony PCR.

### 2.4. Chemical toxicity tests and flask fermentation experiments

Yeast cultures were pre-cultured in YPD medium for 24 h at 250 rpm, then centrifuged at 3,134 ×g for 5 min and washed with distilled water. The cell concentration was adjusted to an optical density at 600 nm (OD_600_) of 0.1 or 1.0 (OD_600_ of 1.0 = 0.6 g of dried cell weight (DCW)/L). For toxicity testing, the growth of *S. cerevisiae* D452-2 and *I. orientalis* NRRL Y-27441 was monitored at OD_600_ using a Synergy H1 microplate reader (Biotek, USA). Pre-cultured yeast cells were inoculated in 200 μL synthetic complete (SC) medium (6.7 g/L yeast nitrogen base and 0.79 g/L complete supplement mixture) with 40 g/L glucose (SCD), supplemented with acetate (0, 2.5, 5, 7.5, 10, 12.5, 15, and 20 g/L) or 3-HP (0, 10, 20, 30, 40, 50, 75, and 100 g/L) at an initial OD_600_ of 0.1, and incubated at 30 □ for 72 h. For flask fermentation, pre-cultured yeast cells were inoculated at OD_600_ 1.0 into 25 mL YP medium containing 40 g/L xylose (YPX) and varying acetate concentrations (0, 2.5, 5, 7.5, 10, 12.5, and 15 g/L) in 125-mL Erlenmeyer flasks. Cultures were incubated at 30 □ and 250 rpm for 120 h.

### 2.5. Industrial raw hemp stalk fermentation

Industrial raw hemp stalks used in this study were harvested from the Throckmorton Purdue Agricultural Center (8343 US-231, Lafayette, IN 47909, USA) (Liu & Beckerman, 2022). The stalks were dried at 60 □ for 24 h, then milled to 50-300 mesh using an Electric Grinder (MFJ-1000, Marada). For dilute acid pretreatment, a 10% (w/v) hemp stalk powder slurry in 1% (w/v) H_2_SO_4_ was treated in a laboratory steam sterilizer (Amsco® Lab 250, Steris, Ireland) at 121 □ for 30 min. The resulting hydrolysate was neutralized to pH 5.5 using 7.5 M NaOH.

For separate hydrolysis and fermentation (SHF), the neutralized hydrolysate (10% w/v) was treated with 30 units/g biomass of cellulase (Celluclast® 1.5L, Novozymes, Bagsvaerd, Denmark) and 30 FBGU/g biomass of hemicellulase (Viscozyme® L, Novozymes, Bagsvaerd, Denmark) at 50 □ for 72 h. Yeast cells (0.6 g DCW/L) and YP medium were then added. For simultaneous saccharification and fermentation (SSF), the neutralized hydrolysate (10% w/v) was inoculated with yeast cells (0.6 g DCW/L), YP medium, cellulase (30 units/g biomass), and hemicellulase (30 FBGU/g biomass). Fermentations were performed at 30 □ and 250 rpm for 96 h.

### 2.6. Analytical methods

Cell density was measured at OD_600_ using a BioMate 160 UV-Vis spectrophotometer (Thermo Scientific, USA). Extracellular metabolites, including glucose, xylose, acetate, 3-HP, and ethanol, were quantified by high-performance liquid chromatography (HPLC) as previously described (Jeong et al., 2024). The column temperature for glucose, xylose, acetate, and ethanol analysis was maintained at 50 □, while for glycerol and 3-HP, it was maintained at 14 □.

Intracellular metabolites were analyzed by gas chromatography-tandem mass spectrometry (GC-MS/MS) following previously described methods (Jeong et al., 2020a) with minor modifications. Pre-cultured *I. orientalis* stains (IoDY01 and IoDY01H) were inoculated at an OD_600_ of 0.1 into YP medium containing 40 g/L xylose with or without 5 g/L acetate. Cultures were harvested during the exponential phase for intracellular metabolite analysis. Intracellular metabolite extraction and derivatization were performed as described (Jeong et al., 2020a). GC-MS/MS analysis was conducted using a Thermo scientific TRACE 1310 GC coupled with a TSQ 8000 MS and TriPlus™ RSH autosampler, with a TraceGOLD TG-5MS column (15 m × 0.25 mm with a 0.25-μm film thickness; Thermo Scientific). One microliter of sample was injected in split mode (51:1) using helium as the carrier gas (1.0 mL/min). The oven temperature program was as follows: 50 □ for 1 min, then ramped at 20 □/min to 330 □, where it was maintained for 5 min. The ion source and transfer line temperatures were set to 250 □ and 290 □, respectively. Ionization was performed using electron impact at 70 eV, and the detector was operated in scan mode with a mass range of 50–550 m/z. Raw data were processed using MS-DIAL software (Tsugawa et al., 2015) (version 5.3) with the Fiehn RI library for mass spectra alignment and metabolite identification. Intracellular 3-HP was quantified by GC-MS/MS.

Chemical composition of the industrial raw hemp stalk (cellulose, hemicellulose, extractives, and lignin) was analyzed using previously described methods (Liu & Beckerman, 2022).

### 2.7. Statistical analysis

Data were presented as the mean ± standard deviation of three biological replicates. Statistical analysis was performed using Microsoft Excel (Office 365) and SPSS version 25 (IBM, Armonk, NY, USA). One-way ANOVA was applied, with post hoc comparisons conducted using Tukey’s HSD test. Statistical significance was defined as *p* < 0.05. Metabolomic data from GC-MS/MS analysis were further processed and analyzed using MetaboAnalyst (version 6.0; https://www.metaboanalyst.ca).

## 3. Results and discussion

### 3.1. Tolerance to acetate and 3-HP in *I. orientalis*

To assess the tolerance of *I. orientalis* for 3-HP production from lignocellulosic biomass with high acetate content (e.g., hemp biomass), its tolerance to acetate and 3-HP was compared with that of the model yeast *S. cerevisiae* (Fig. 1). Under acetate stress, *S. cerevisiae* and *I. orientalis* grew up to 2.5 and 5.0 g/L acetate, respectively. Notably, *I. orientalis* growth was unaffected at 5.0 g/L acetate but completely inhibited at 10 g/L acetate (Fig. 1A). Under 3-HP stress, *S. cerevisiae* growth was inhibited at varying concentrations, while *I. orientalis* growth was significantly reduced only at 75 g/L 3-HP (Fig. 1B). These results confirm that *I. orientalis* strain has higher tolerance to acetate compared to *S. cerevisiae*, consistent with previous reports (Seong et al., 2017).

**Fig. 1.**
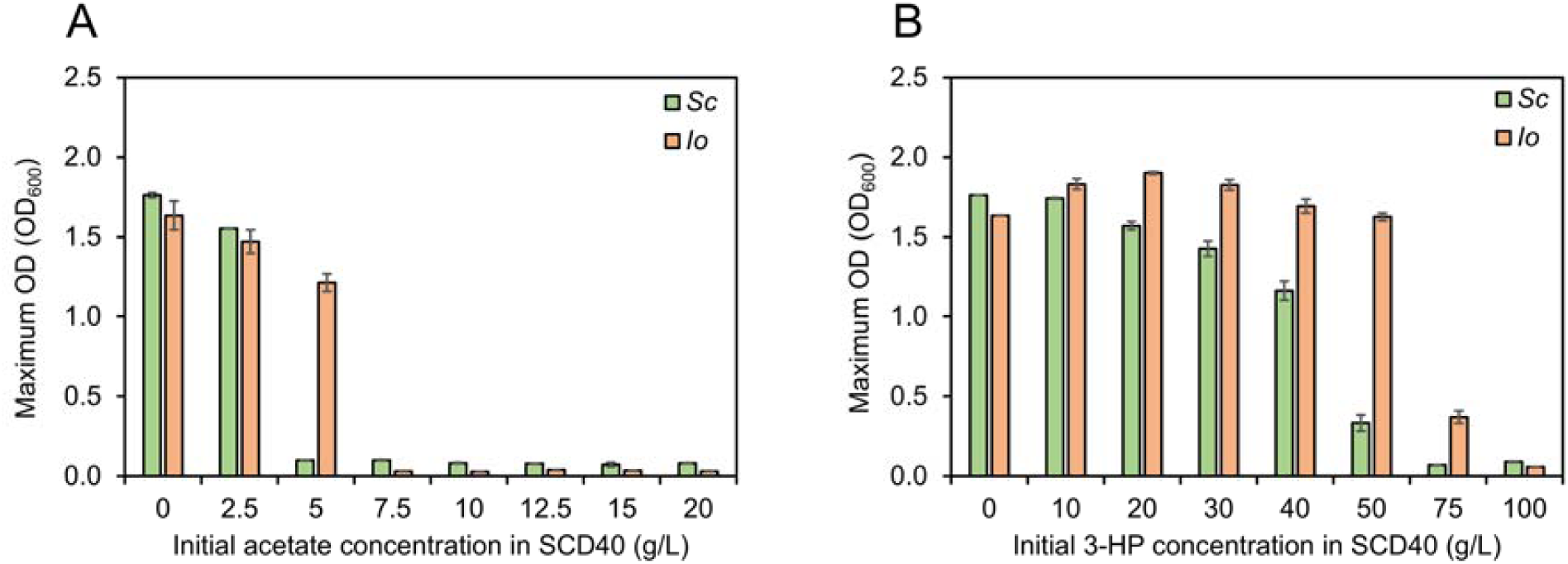
Tolerance of *S. cerevisiae* and *I. orientalis* to acetate and 3-hydroxypropionic acid (3-HP) Wild-type *S. cerevisiae* D452-2 (*Sc*) and wild-type *I. orientalis* NRRL Y-27441 (*Io*) strains were grown in SC medium containing 40 g/L glucose (SCD40) with increasing concentrations of acetate (A; 0, 2.5, 5, 7.5, 10, 12.5, 15, and 20 g/L) or 3-HP (B; 0, 10, 20, 30, 40, 50, 75, and 100 g/L). Growth was monitored over 72 h using a microplate reader. Data represent biological triplicates; error bars indicate standard deviation.

### 3.2. Construction of engineered *I. orientalis* capable of 3-HP production from xylose

Diversification of selection markers enhances genetic engineering flexibility, supports industrial microorganism utilization, and facilitates multiple genetic modifications (Fels et al., 2020; Jillette et al., 2019; Tan et al., 2025). However, the genetic engineering of *I. orientalis* predominantly relies on auxotrophic markers such as *URA3* (Kitagawa et al., 2010; Xiao et al., 2014) and *LEU2* (Wu et al., 2023). Although antibiotic resistance markers such as nourseothricin sulfate (Lee et al., 2022b) and zeocin (Zhang et al., 2023) have been used recently, there remains a need for broader marker diversity.

To advance Cas9-based genome editing in *I. orientalis*, we constructed a Cas9 plasmid incorporating novel antibiotic resistance markers and expression cassettes (Fig. S1A). First, we evaluated potential antibiotic markers in five wild-type *I. orientalis* strains (SD108, NRRL Y-27441, ATCC MBY 1358, ATCC 24210, and NRRL YB-373) cultured on YPD agar containing G418 sulfate (300, 350, 400, 500, and 600 μg/mL) or hygromycin B (300, 350, 400, 500, and 600 μg/mL). Growth of four strains was inhibited at 500 μg/mL G418 sulfate, while the ATCC MBY 1358 strain tolerated concentrations up to 600 μg/mL (Fig. S2B). Conversely, all strains were inhibited at 300 μg/mL hygromycin B (Fig. S1C). These findings align with prior reports indicating variable resistance to G418 and greater uniform sensitivity to hygromycin B (Pyne et al., 2023). Thus, although G418 sulfate can serve as a selection marker, its high required concentration reduces cost-effectiveness. In contrast, hygromycin B offers a more practical selective marker for *I. orientalis*. To demonstrate its utility, we replaced the nourseothricin sulfate resistance (*Nat*) marker in the pCast plasmid with a hygromycin B resistance marker (*hph*) using gibson assembly, creating the pCastH plasmid (Fig. S1A). The Y-27441 strain harboring pCastH grew in YPD liquid medium containing 300 μg/mL hygromycin B (Fig. S2D).

For 3-HP production from xylose, heterologous genes of the xylose oxidoreductase pathway and the β-alanine-based 3-HP pathway were integrated into the *I. orientalis* Y-27441 genome using Cas9-based genome editing with a hygromycin resistance marker (Fig. 2). First, we introduced *XYL1* (xylose reductase), *XYL2* (xylitol dehydrogenase), and *XYL3* (xylulokinase) from *Scheffersomyces stipitis* to enhance xylose consumption (Jeong et al., 2020a; Lee et al., 2022b), leading to the development of the IoDY01 strain. Next, for 3-HP biosynthesis, we introduced *TcPAND* (aspartate decarboxylase) from *Tribolium castaneum*, *BcBAPAT* (β-alanine-pyruvate aminotransferase) from *Bacillus cereus*, and *EcydfG* (3-hydroxypropionate dehydrogenase) from *E. coli* (Kildegaard et al., 2015) into the genome of IoDY01. Each gene was placed between the corresponding promoter and terminator of the expression cassettes, generating the engineered IoDY01H strain.

**Fig. 2.**
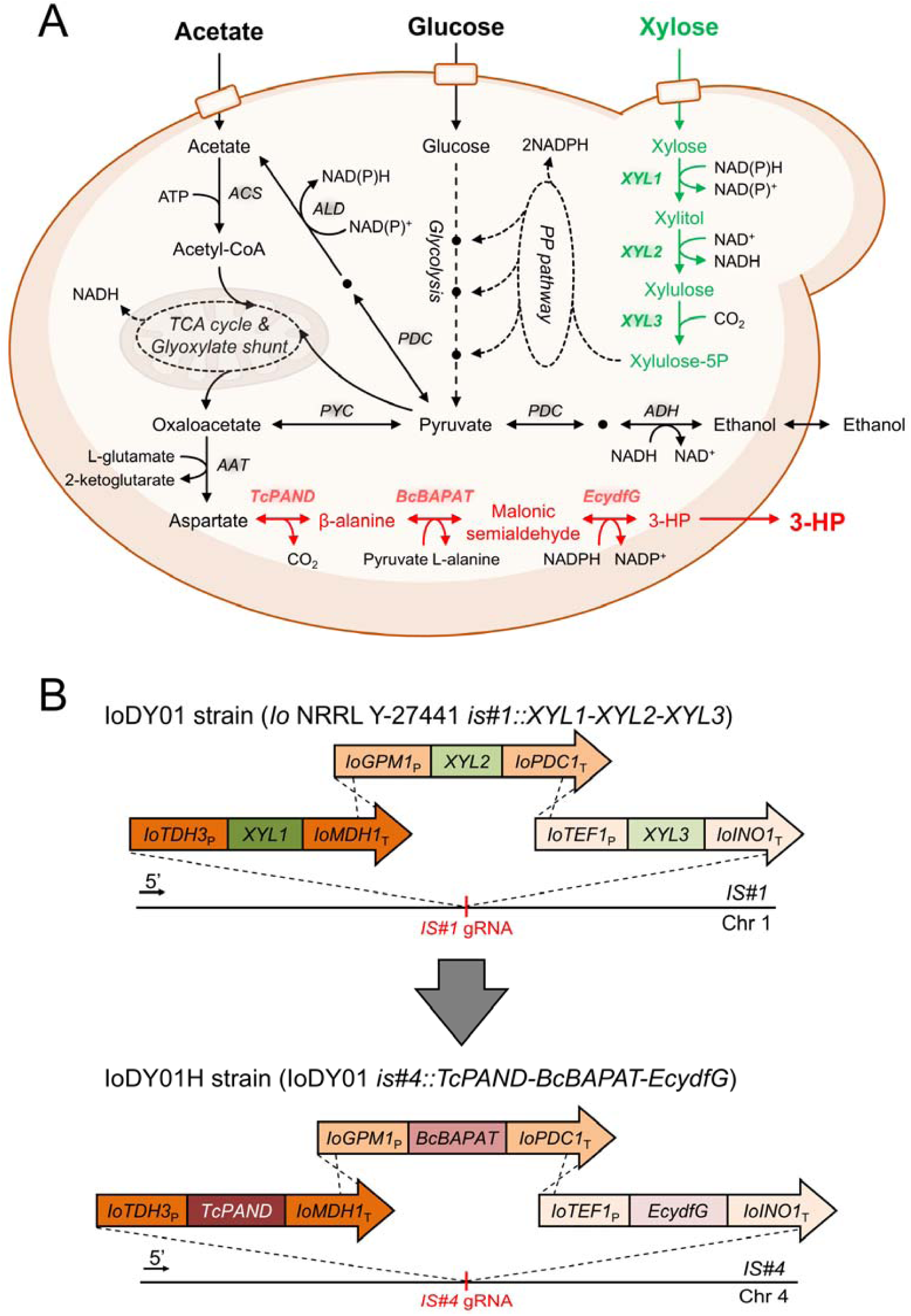
Schematic diagram of 3-HP production from various substrates (xylose, glucose, and acetate) in an engineered *I. orientalis* strain. (A) Metabolic pathways introduced into *I. orientalis*, including the heterologous xylose oxidoreductase pathway genes (*XYL1*, *XYL2*, and *XYL3*) and β-alanine-based 3-HP biosynthetic pathway genes (*TcPAND*, *BcBAPAT*, and *EcydfG*). (B) Metabolic engineering strategy for introducing xylose and 3-HP pathway expression cassettes into *I. orientalis*. Five heterologous genes (*XYL1*, *XYL2*, *XYL3*, *TcPAND*, and *BcBAPAT*) were codon optimized. *ACS*, acetyl-CoA synthetase; *ALD*, aldehyde dehydrogenase; *PDC*, pyruvate decarboxylase; *PYC*, pyruvate carboxylase; *ADH*, alcohol dehydrogenase; *AAT*, aspartate aminotransferase; *XYL1*, xylose reductase derived from *Scheffersomyces stipitis*; *XYL2*, xylitol dehydrogenase derived from *Scheffersomyces stipitis*; *XYL3*, xylulokinase derived from *Scheffersomyces stipitis*; *TcPAND*, aspartate-1-decarboxylase derived from *Tribolium castaneum*; *BcBAPAT*, β-alanine-pyruvate aminotransferase derived from *Bacillus cereus*; *EcydfG*, 3-hydroxypropionate dehydrogenase derived from *Escherichia coli*.

### 3.3. 3-HP production from mixed carbon sources of xylose and acetate in the engineered *I. orientalis* strain

The metabolomic profiles of the engineered *I. orientalis* strains were analyzed to assess 3-HP production from xylose. The IoDY01 strain fully consumed xylose as the sole carbon source within 36 hours, producing 8.8 g/L xylitol and 4.0 g/L ethanol (Fig. 3A), similar to the previously reported IO21X strain (Lee et al., 2022b). In contrast, the IoDY01H strain expressing the 3-HP pathway genes (*TcPAND*, *BcBAPAT*, and *EcydfG*) produced 4.8 g/L of 3-HP but exhibited 26.8% and 34.4% reductions in cell growth and ethanol accumulation, respectively, likely due to competition for pyruvate-derived precursors (Fig. 3B).

**Fig. 3.**
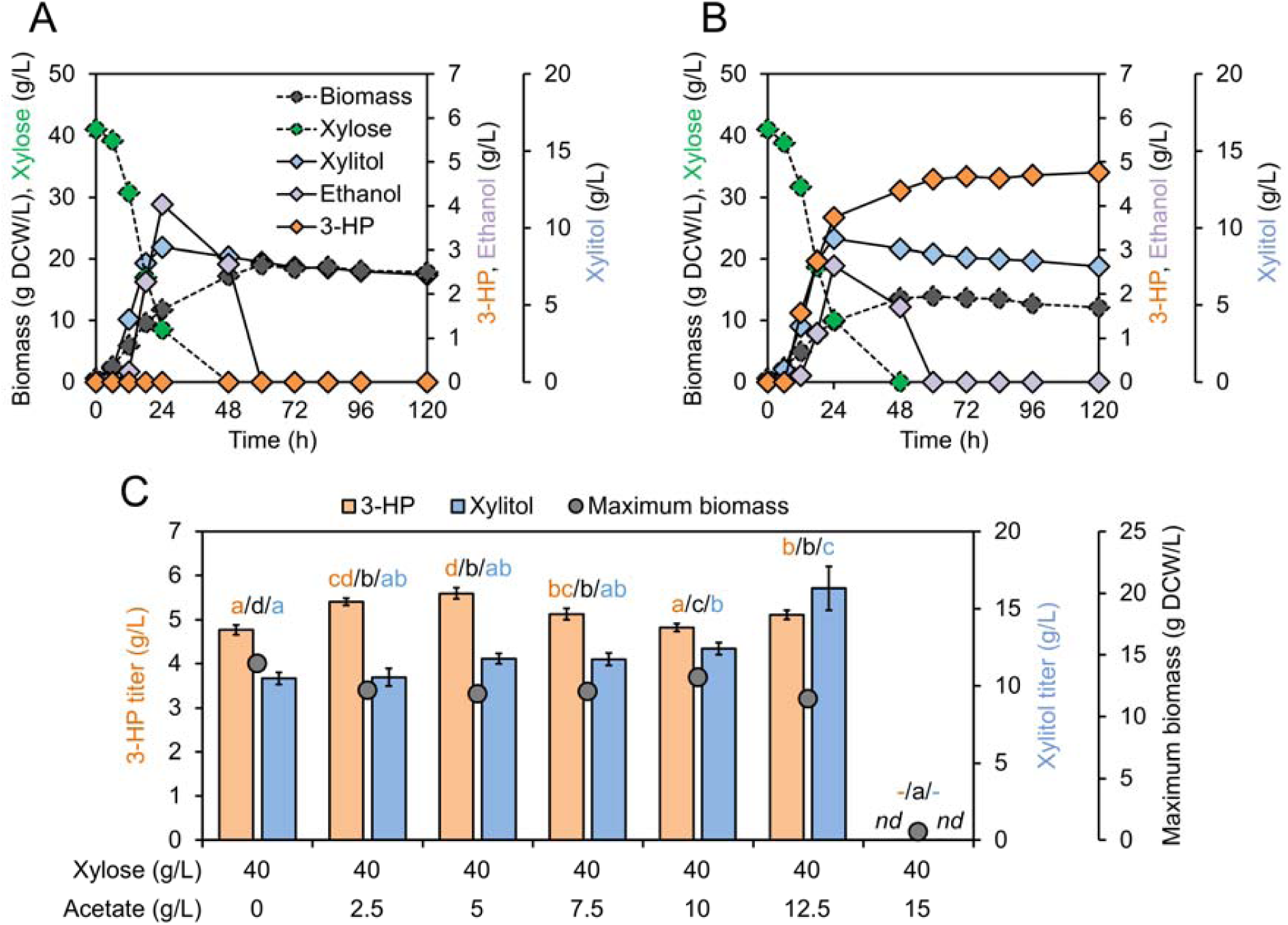
Production of 3-HP from xylose or mixed carbon sources of xylose and acetate in engineered *I. orientalis* strains. (A and B) Xylose fermentation profiles of engineered strains IoDY01 (A) and IoDY01H (B). (C) Comparison of maximum biomass (black), 3-HP production (light orange), and xylitol production (light blue) over 120 h from mixed carbon sources of xylose and acetic acid using the engineered IoDY01H strain. Strains were cultured in YP medium containing (A and B) 40 g/L xylose or (C) 40 g/L xylose with varying concentrations of acetate (0, 2.5, 5, 7.5, 10, 12.5, and 15 g/L) at 30 □ and 250 rpm for 120 h with an initial cell density of 0.6 g DCW/L. All experiments were performed in biological triplicate; error bars represent standard deviation. Different letters indicate statistically significant differences (Tukey’s HSD tests, *p* < 0.05). *nd*, not detected.

To examine the impact of acetate supplementation on 3-HP production from xylose, fermentation was performed with increasing acetate concentrations (0, 2.5, 5, 7.5, 10, 12.5, and 15 g/L) using the IoDY01H strain (Fig. 3). Acetate addition reduced maximum cell growth, with complete inhibition of xylose consumption observed at 15 g/L acetate (Fig. 3C and S3). However, co-consumption of xylose and acetate enhanced 3-HP production, reaching a maximum titer of 5.6 g/L at 5 g/L acetate, while xylitol accumulation increased by up to 46.5% with rising acetate concentrations (Fig. 3C). These results suggest that carbon flux from acetate metabolism was preferentially directed toward intermediate metabolites supporting the biosynthesis of 3-HP and xylitol, rather than biomass accumulation. Despite elevated acetate availability, only a modest increase in 3-HP was observed, while xylitol accumulation was substantially enhanced. This suggests that acetate metabolism inhibits NAD^+^-dependent xylitol dehydrogenase (Xyl2), a key enzyme in the xylose assimilation pathway, in an acetate-dependent manner. A previous study reported that acetate metabolism in an engineered *Kluyveromyces marxianus* strain activated the glyoxylate shunt during xylose fermentation, increasing energy production and NADH cofactor biosynthesis. This, in turn, suppressed the activity of NAD^+^-dependent xylitol dehydrogenase (Xyl2), resulting in elevated xylitol accumulation (Du et al., 2024). These findings indicate that xylose served as a limiting energy source for 3-HP production under high acetate concentrations.

### 3.4. Intracellular metabolite profiles during xylose fermentation with acetate supplementation

To evaluate intracellular metabolite changes and identify bottlenecks during 3-HP production, four experimental conditions were tested: Con_X (IoDY01 with xylose), MT_X (IoDY01H with xylose), Con_XAa (IoDY01 with xylose and acetate), and MT_XAa (IoDY01H with xylose and acetate) (Fig. 4). Principal component analysis (PCA) revealed that PC1 captured metabolomic shifts driven by acetate supplementation, whereas PC2 accounted for differences associated with the 3-HP pathway (Fig. 4B).

**Fig. 4.**
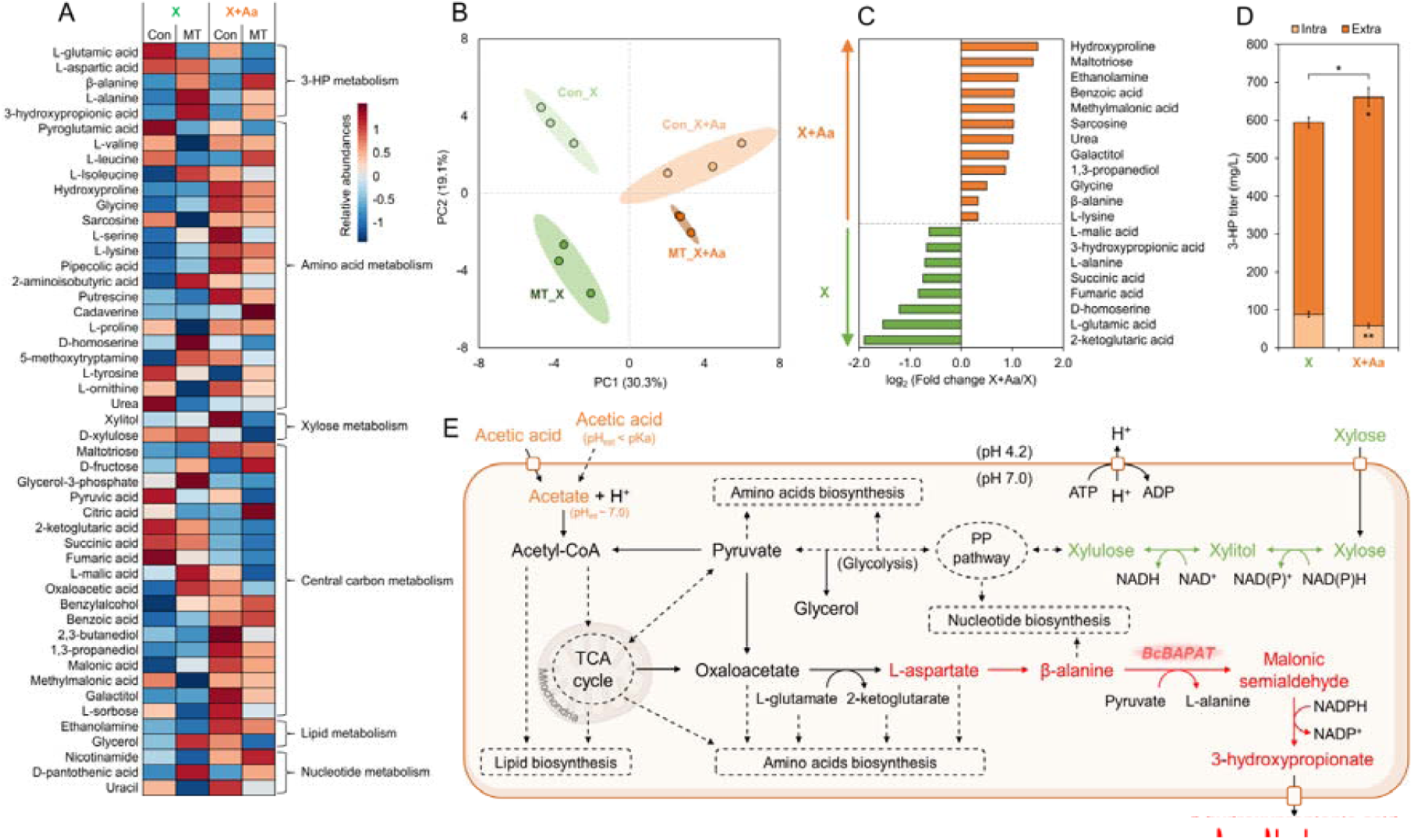
Comparative metabolomic analysis of IoDY01 and IoDY01H strains during xylose fermentation without or with acetate supplementation. (A) Heatmap of 49 intracellular metabolites detected by GC-MS/MS in IoDY01 (Con) and IoDY01H (MT) strains cultured with xylose alone (X) or xylose plus acetate (X+Aa). Data were analyzed using Pearson correlation and visualized with MetaboAnalyst 6.0. Metabolites were grouped by KEGG pathways. Relative abundance is represented from low (blue) to intermediate (white) to high (red). (B) Principal component analysis (PCA) score plot (F = 36.068; R^2^ = 0.93116; *p* = 0.001) of intracellular metabolites from Con_X (light green), Con_X+Aa (light orange), MT_X (green), and Con_X+Aa (orange). Each dot represents a biological replicate, and ellipses indicate 95% confidence intervals. (C) Significantly altered intracellular metabolites in IoDY01H in response to acetate supplementation (Student’s t-test; *p* < 0.05). Metabolites enriched with acetate addition are shown in orange (positive fold change), and those without acetate in green (negative fold change). (D) Titers of intracellular (Intra) and extracellular (Extra) 3-HP in IoDY01H during xylose fermentation with acetate. Statistical significance was determined by two-tailed unpaired Student’s t-test (*, *p* < 0.05; **, *p* < 0.01). (E) Schematic representation of the metabolic pathway for 3-HP biosynthesis from xylose and acetate. All strains were cultured in YP medium containing 40 g/L xylose with or without 5 g/L acetate at 30 □ and 250 rpm, starting from an initial OD_600_ of 0.1. Cells were harvested during the exponential phase for intracellular metabolite analysis. All experiments were conducted in biological triplicate.

A total of 49 intracellular metabolites were identified and categorized into amino acid metabolism, xylose metabolism, glycolysis, central carbon metabolism—comprising glycolysis and the tricarboxylic acid (TCA) cycle, lipid metabolism, nucleotide metabolism, and 3-HP metabolism, as classified by the KEGG pathway (Kanehisa & Goto, 2000) (Fig. 4A). Independent of the 3-HP pathway, acetate supplementation decreased the levels of intermediate metabolites in the TCA cycle (pyruvic acid, 2-ketoglutaric acid, succinic acid, fumaric acid, L-malic acid, and oxaloacetic acid), while increasing intermediates associated with lipid, nucleotide, and amino acid metabolism (Fig. 4A). Previous transcriptomic analyses reported that genes involved in energy metabolism, including the TCA cycle, were down-regulated in *S. cerevisiae* under organic acid stress but up-regulated in acid-tolerant yeasts such as *I. orientalis* (Li et al., 2022; Li et al., 2020). Also, transcriptomic and metabolomic data integration studies support the decoupling of transcriptome and metabolome profiles, with significant inverse correlations reported between enzyme abundance and substrate metabolite concentrations in yeast central carbon metabolism (Cavill et al., 2016; Fendt et al., 2010). These observations suggest that during xylose fermentation, acetate supplementation upregulates the TCA cycle and redirects intermediates toward amino acids and lipid biosynthesis. This reinforces the inverse relationship between transcript levels and metabolite concentrations (Fig. 4A).

Expression of the β-alanine-based 3-HP biosynthetic pathway led to increased accumulation of β-alanine, L-alanine, and 3-HP, indicating a redirection of metabolic flow toward 3-HP production (Fig. 4A). However, acetate supplementation reduced levels of L-alanine and 3-HP compared to xylose-only fermentation (Fig. 4C), suggesting that β-alanine-pyruvate aminotransferase, which catalyzes the conversion of β-alanine and pyruvate into malonic semialdehyde and L-alanine, may serve as a key bottleneck in 3-HP production. The optimal pH of β-alanine-pyruvate aminotransferase derived from *B. cereus* (BcBAPAT) is 10.0, with residual activity of 40% under neutral pH conditions (Nakano et al., 1977). Therefore, engineering this enzyme to improve its efficiency under acidic conditions is necessary. Furthermore, co-metabolism of xylose and acetate in IoDY01H decreased the concentrations of L-glutamic acid, 2-ketoglutaric acid, and L-aspartic acid, but β-alanine levels increased, compared to xylose only fermentation (Fig. 4C). Since these reduced intermediate metabolites serve as precursors for amino acid biosynthesis (Takagi, 2019), the results suggest that acetate redirects metabolic flux toward amino acid biosynthesis rather than respiration, ultimately enhancing 3-HP biosynthesis.

Acetic acid supplementation has been reported to upregulate membrane proteins, transporters, and ATPases (Dubinkina et al., 2024) (Fig. 4E), and the activation of the pH homeostasis mechanism under acid stress contributes to the regulation of citric acid production and excretion (Ochoa-Estopier & Guillouet, 2014). Consistent with these findings, the addition of acetate significantly altered 3-HP distribution, decreasing the intracellular 3-HP titer from 88.3 mg/L to 58.5 mg/L, while increasing the extracellular 3-HP titer from 505.7 mg/L to 602.3 mg/L. This shift resulted in an overall increase in total 3-HP titer from 594.0 mg/L to 660.9 mg/L (Fig. 4D). This regulatory effect may indirectly influence 3-HP secretion. However, in an acidified culture medium (pH < 3-HP pKa of 4.51), 3-HP may be reabsorbed, reducing overall production yield (Qin et al., 2024). The transient nature of pH homeostasis suggests that additional engineering of transporters and permeases will be required for sustained 3-HP production.

### 3.5. 3-HP production from the industrial hemp stalk fermentation

To evaluate the optimal approach for 3-HP production from acetate-rich lignocellulosic biomass, we compared simultaneous saccharification and fermentation (SSF) and separate hydrolysis and fermentation (SHF) using the engineered IoDY01H strain and hemp stalk biomass as a substrate. The raw help stalk contained 59.42□±□0.86% cellulose, 18.86□±□0.72% hemicellulose, 2.37□±□0.27% extractives, 0.51□±□0.09% acid-soluble lignin, and 19.69□±□0.42% acid-insoluble lignin. This higher lignin content, compared to previous reports, likely reflects variations in plant organ composition (Väisänen et al., 2019).

SSF offers faster cell growth and higher productivity due to low initial inhibitor levels but is constrained by low hydrolysis efficiency due to reduced temperatures required for microbial fermentation (Choudhary et al., 2016). In contrast, SHF enables higher initial sugar yields, albeit at the expense of increased acetate accumulation (Guo et al., 2023). In the SHF process, 10% (w/v) hemp stalk hydrolysate contained 23.2 ± 0.7 g/L glucose, 16.8 ± 0.9 g/L xylose, and 4.3 ± 0.5 g/L acetate, while the SSF process yielded 18.1 ± 0.1 g/L glucose, 15.4 ± 0.0 g/L xylose, and 3.1 ± 0.0 g/L acetate. Despite the higher initial acetate levels in SHF, the IoDY01H strain achieved a higher 3-HP titer (8.7 ± 0.4 g/L) than SSF (7.7 ± 0.0 g/L), corresponding to a 3-HP yield of 0.20 g/g (Fig. 5). This indicates that *I. orientalis* effectively mitigates acetate inhibition and simultaneously consumes glucose, xylose, and acetate, thereby maintaining consistent 3-HP productivity. This study demonstrates that acid-tolerant *I. orientalis* enables feasible 3-HP production and has strong potential for efficient lignocellulosic biomass bioconversion with minimal genetic manipulation (Tan et al., 2025; Wang et al., 2025).

**Fig. 5.**
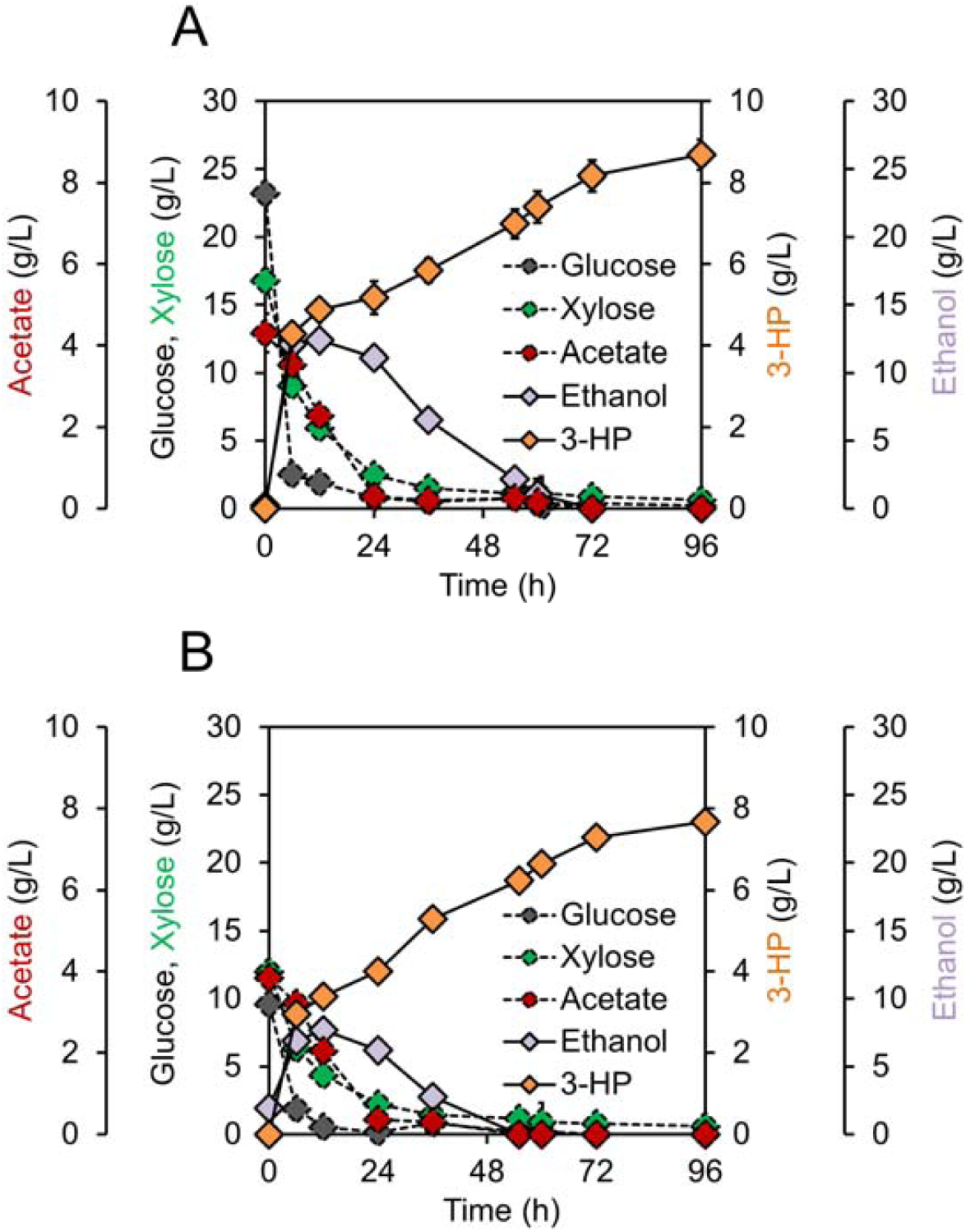
Comparison of SHF and SSF processes for 3-HP production from acetate-rich hemp stalk hydrolysate using the IoDY01H strain. (A) Fermentation profiles of separate hydrolysis and fermentation (SHF). Neutralized hemp stalk hydrolysate (10% w/v) was pretreated with cellulase (30 units/g biomass) and hemicellulase (30 FBGU/g biomass) at 50 □ for 72 h. Subsequently, pre-cultured IoDY01H cells (0.6 g DCW/L), yeast extract (10 g/L), and peptone (20 g/L) were added. (B) Fermentation profiles of simultaneous saccharification and fermentation (SSF). Neutralized hemp stalk hydrolysate (10% w/v) was co-inoculated with IoDY01H cells (0.6 g DCW/L), yeast extract (10 g/L), peptone (20 g/L), cellulase (30 units/g biomass), and hemicellulase (30 FBGU/g biomass). All fermentations were performed in triplicate at 30 □ and under aerobic conditions (250 rpm).

## 4. Conclusion

This study highlights the potential of an acid-tolerant *I. orientalis* strain for efficient 3-HP production from acetate-rich biomass. A Cas9-based genetic toolkit carrying a hygromycin B antibiotic marker was developed, enabling the construction of the IoDY01H strain, which achieved a 3-HP titer of 8.7 g/L using the SHF process from hemp stalk hydrolysate. This yield was higher than that achieved in the SSF process. Metabolomic analysis revealed that acetate metabolism during xylose fermentation facilitates the accumulation of key intermediates for 3-HP biosynthesis. These results underscore the viability of *I. orientalis* as a robust platform for converting acetate, a common lignocellulosic fermentation inhibitor, into high-value chemicals.

## CRediT authorship contribution statement

**Deokyeol Jeong**: Conceptualization, Data curation, Formal analysis, Investigation, Methodology, Software, Project administration, Writing – original draft, Writing – review & editing, Visualization. **Dahye Lee**: Data curation, Formal analysis, Writing – review & editing, Visualization. **Junli Liu**: Data curation, Formal analysis, Resources. **Soo Rin Kim**: Resources, Writing – review & editing. **Yong-Su Jin**: Resources, Writing – review & editing. **Jikai Zhao**: Writing – review & editing. **Eun Joong Oh**: Conceptualization, Funding acquisition, Project administration, Resources, Supervision, Writing – original draft, Writing – review & editing.

## Acknowledgments

This work was supported by the U.S. Department of Agriculture (Agriculture and Food Research Initiative, Biorefining and Biomanufacturing Program, Grant Number 2022-67022-37609). We thank Dr. Nurzhan Kuanyshev at the University of Illinois at Urbana–Champaign for providing information on the pCast plasmid.

## Appendix A Supplementary data

E-supplementary data for this work can be found in the online version of this paper.

## References

Abbott, D.A., Knijnenburg, T.A., De Poorter, L.M., Reinders, M.J., Pronk, J.T., Van Maris, A.J. 2007. Generic and specific transcriptional responses to different weak organic acids in anaerobic chemostat cultures of *Saccharomyces cerevisiae*. FEMS Yeast Research, 7(6), 819–833.

Barnhart, M., Negrete-Raymond, A., Frias, J., Barbier, G., Catlett, M. 2017. 3-hydroxypropionic acid production by recombinant yeasts expressing an insect aspartate 1-decarboxylase, Google Patents.

Benatuil, L., Perez, J.M., Belk, J., Hsieh, C.-M. 2010. An improved yeast transformation method for the generation of very large human antibody libraries. Protein Engineering, Design & Selection, 23(4), 155–159.

Bozell, J.J., Petersen, G.R. 2010. Technology development for the production of biobased products from biorefinery carbohydrates—the US Department of Energy’s “Top 10” revisited. Green Chemistry, 12(4), 539–554.

Cavill, R., Jennen, D., Kleinjans, J., Briedé, J.J. 2016. Transcriptomic and metabolomic data integration. Briefings in Bioinformatics, 17(5), 891–901.

Choi, S., Song, C.W., Shin, J.H., Lee, S.Y. 2015. Biorefineries for the production of top building block chemicals and their derivatives. Metabolic Engineering, 28, 223–239.

Choi, S.Y., Rhie, M.N., Kim, H.T., Joo, J.C., Cho, I.J., Son, J., Jo, S.Y., Sohn, Y.J., Baritugo, K.-A., Pyo, J. 2020. Metabolic engineering for the synthesis of polyesters: A 100-year journey from polyhydroxyalkanoates to non-natural microbial polyesters. Metabolic Engineering, 58, 47–81.

Choudhary, J., Singh, S., Nain, L. 2016. Thermotolerant fermenting yeasts for simultaneous saccharification fermentation of lignocellulosic biomass. Electronic Journal of Biotechnology, 21, 82–92.

de Fouchécour, F., Sánchez-Castañeda, A.-K., Saulou-Berion, C., Spinnler, H.E. 2018. Process engineering for microbial production of 3-hydroxypropionic acid. Biotechnology Advances, 36(4), 1207–1222.

Du, C., He, Y., Liu, J., Su, L., Li, Y., Yuan, W., Bai, F. 2024. Metabolic coupling of acetate promotes xylose utilization in *Kluyveromyces marxianus*. Chemical Engineering Journal, 484, 149762.

Dubinkina, V., Bhogale, S., Hsieh, P.-H., Dibaeinia, P., Nambiar, A., Maslov, S., Yoshikuni, Y., Sinha, S. 2024. A transcriptomic atlas of acute stress response to low pH in multiple *Issatchenkia orientalis* strains. Microbiology Spectrum, 12(1), e02536–23.

Fels, U., Gevaert, K., Van Damme, P. 2020. Bacterial genetic engineering by means of recombineering for reverse genetics. Frontiers in Microbiology, 11, 548410.

Fendt, S.M., Buescher, J.M., Rudroff, F., Picotti, P., Zamboni, N., Sauer, U. 2010. Tradeoff between enzyme and metabolite efficiency maintains metabolic homeostasis upon perturbations in enzyme capacity. Molecular Systems Biology, 6(1), 356.

Feng, J., Techapun, C., Phimolsiripol, Y., Phongthai, S., Khemacheewakul, J., Taesuwan, S., Mahakuntha, C., Porninta, K., Htike, S.L., Kumar, A. 2023. Utilization of agricultural wastes for co-production of xylitol, ethanol, and phenylacetylcarbinol: A review. Bioresource Technology, 129926.

Ge, H., Zheng, J., Xu, H. 2023. Advances in machine learning for high value-added applications of lignocellulosic biomass. Bioresource Technology, 369, 128481.

Gietz, R.D., Schiestl, R.H. 2007. High-efficiency yeast transformation using the LiAc/SS carrier DNA/PEG method. Nature Protocols, 2(1), 31–34.

Gong, G., Wu, B., Liu, L., Li, J., Zhu, Q., He, M., Hu, G. 2022. Metabolic engineering using acetate as a promising building block for the production of bio□based chemicals. Engineering Microbiology, 2(4), 100036.

Guaragnella, N., Bettiga, M. 2021. Acetic acid stress in budding yeast: From molecular mechanisms to applications. Yeast, 38(7), 391–400.

Guo, H., Zhao, Y., Chang, J.-S., Lee, D.-J. 2023. Enzymes and enzymatic mechanisms in enzymatic degradation of lignocellulosic biomass: a mini-review. Bioresource technology, 367, 128252.

Hosaka, K., Nikawa, J.-i., Kodaki, T., Yamashita, S. 1992. A dominant mutation that alters the regulation of *INO1* expression in *Saccharomyces cerevisiae*. The Journal of Biochemistry, 111(3), 352–358.

Huang, G., Li, J., Lin, J., Duan, C., Yan, G. 2024. Multi-modular metabolic engineering and efflux engineering for enhanced lycopene production in recombinant *Saccharomyces cerevisiae*. Journal of Industrial Microbiology and Biotechnology, 51, kuae015.

Jeong, D., Oh, E.J., Ko, J.K., Nam, J.-O., Park, H.-S., Jin, Y.-S., Lee, E.J., Kim, S.R. 2020a. Metabolic engineering considerations for the heterologous expression of xylose-catabolic pathways in *Saccharomyces cerevisiae*. PLoS One, 15(7), e0236294.

Jeong, D., Park, S., Evelina, G., Kim, S., Park, H., Lee, J.M., Kim, S.-K., Kim, I.J., Oh, E.J., Kim, S.R. 2024. Bioconversion of citrus waste into mucic acid by xylose-fermenting *Saccharomyces cerevisiae*. Bioresource Technology, 393, 130158.

Jeong, D., Ye, S., Park, H., Kim, S.R. 2020b. Simultaneous fermentation of galacturonic acid and five-carbon sugars by engineered *Saccharomyces cerevisiae*. Bioresource Technology, 295, 122259.

Jillette, N., Du, M., Zhu, J.J., Cardoz, P., Cheng, A.W. 2019. Split selectable markers. Nature Communications, 10(1), 4968.

Kanehisa, M., Goto, S. 2000. KEGG: kyoto encyclopedia of genes and genomes. Nucleic Acids Research, 28(1), 27–30.

Kildegaard, K.R., Wang, Z., Chen, Y., Nielsen, J., Borodina, I. 2015. Production of 3-hydroxypropionic acid from glucose and xylose by metabolically engineered *Saccharomyces cerevisiae*. Metabolic Engineering Communications, 2, 132–136.

Kim, S., Jeong, D., Jang, B., Park, S., Oh, E.J., Kim, I.J., Kim, S.R. 2024. Coupled engineering strategy of *CYB2* deletion and *ACS1* overexpression improves cellulosic lactic acid production by *Saccharomyces cerevisiae*. Biomass and Bioenergy, 185, 107249.

Kitagawa, T., Tokuhiro, K., Sugiyama, H., Kohda, K., Isono, N., Hisamatsu, M., Takahashi, H., Imaeda, T. 2010. Construction of a β-glucosidase expression system using the multistress-tolerant yeast *Issatchenkia orientalis*. Applied Microbiology and Biotechnology, 87, 1841–1853.

Lee, Y.-G., Ju, Y., Sun, L., Park, S., Jin, Y.-S., Kim, S.R. 2022a. Acetate-rich cellulosic hydrolysates and their bioconversion using yeasts. Biotechnology and Bioprocess Engineering, 27(6), 890–899.

Lee, Y.-G., Kang, N.K., Kim, C., Tran, V.G., Cao, M., Yoshikuni, Y., Zhao, H., Jin, Y.-S. 2024. Self-Buffering system for Cost-Effective production of lactic acid from glucose and xylose using Acid-Tolerant *Issatchenkia orientalis*. Bioresource Technology, 399, 130641.

Lee, Y.-G., Kim, C., Kuanyshev, N., Kang, N.K., Fatma, Z., Wu, Z.-Y., Cheng, M.-H., Singh, V., Yoshikuni, Y., Zhao, H. 2022b. Cas9-based metabolic engineering of *Issatchenkia orientalis* for enhanced utilization of cellulosic hydrolysates. Journal of Agricultural and Food Chemistry, 70(38), 12085–12094.

Li, M., Chu, Y., Dong, X., Ji, H. 2024. General mechanisms of weak acid-tolerance and current strategies for the development of tolerant yeasts. World Journal of Microbiology and Biotechnology, 40(2), 49.

Li, Y., Li, Y., Li, R., Liu, L., Miao, Y., Weng, P., Wu, Z. 2022. Metabolic changes of *Issatchenkia orientalis* under acetic acid stress by transcriptome profile using RNA-sequencing. International Microbiology, 1–10.

Li, Y., Wu, Z., Li, R., Miao, Y., Weng, P., Wang, L. 2020. Integrated transcriptomic and proteomic analysis of the acetic acid stress in *Issatchenkia orientalis*. Journal of Food Biochemistry, 44(6), e13203.

Liang, J., Liu, S., Du, Z., Zhang, R., Lv, L., Sun, L., Nabi, M., Zhang, G., Zhang, P. 2024. Recent advances in methane and hydrogen production from lignocellulosic degradation with anaerobic fungi. Bioresource Technology, 131544.

Liu, J., Beckerman, J. 2022. Application of sustainable biosorbents from hemp for remediation copper (II)-containing wastewater. Journal of Environmental Chemical Engineering, 10(3), 107494.

Mutyala, S., Kim, J.R. 2022. Recent advances and challenges in the bioconversion of acetate to value-added chemicals. Bioresource Technology, 364, 128064.

Nakano, Y., Tokunaga, H., Kitaoka, S. 1977. Two ω-amino acid transaminases from *Bacillus cereus*. The Journal of Biochemistry, 81(5), 1375–1381.

Ochoa-Estopier, A., Guillouet, S.E. 2014. D-stat culture for studying the metabolic shifts from oxidative metabolism to lipid accumulation and citric acid production in *Yarrowia lipolytica*. Journal of Biotechnology, 170, 35–41.

Pyne, M.E., Bagley, J.A., Narcross, L., Kevvai, K., Exley, K., Davies, M., Wang, Q., Whiteway, M., Martin, V.J. 2023. Screening non-conventional yeasts for acid tolerance and engineering *Pichia occidentalis* for production of muconic acid. Nature Communications, 14(1), 5294.

Qin, N., Li, L., Wan, X., Ji, X., Chen, Y., Li, C., Liu, P., Zhang, Y., Yang, W., Jiang, J. 2024. Increased CO_2_ fixation enables high carbon-yield production of 3-hydroxypropionic acid in yeast. Nature Communications, 15(1), 1591.

Seong, Y.J., Lee, H.J., Lee, J.E., Kim, S., Lee, D.Y., Kim, K.H., Park, Y.C. 2017. Physiological and metabolomic analysis of *Issatchenkia orientalis* MTY1 with multiple tolerance for cellulosic bioethanol production. Biotechnology Journal, 12(11), 1700110.

Shi, W., Li, J., Chen, Y., Liu, X., Chen, Y., Guo, X., Xiao, D. 2021. Metabolic engineering of *Saccharomyces cerevisiae* for ethyl acetate biosynthesis. ACS Synthetic Biology, 10(3), 495–504.

Sun, L., Lee, J.W., Yook, S., Lane, S., Sun, Z., Kim, S.R., Jin, Y.-S. 2021. Complete and efficient conversion of plant cell wall hemicellulose into high-value bioproducts by engineered yeast. Nature Communications, 12(1), 4975.

Sun, W., Vila-Santa, A., Liu, N., Prozorov, T., Xie, D., Faria, N.T., Ferreira, F.C., Mira, N.P., Shao, Z. 2020. Metabolic engineering of an acid-tolerant yeast strain *Pichia kudriavzevii* for itaconic acid production. Metabolic Engineering Communications, 10, e00124.

Suthers, P.F., Maranas, C.D. 2022. Examining organic acid production potential and growth□coupled strategies in *Issatchenkia orientalis* using constraint□based modeling. Biotechnology Progress, 38(5), e3276.

Takagi, H. 2019. Metabolic regulatory mechanisms and physiological roles of functional amino acids and their applications in yeast. *Bioscience*, Biotechnology, and Biochemistry, 83(8), 1449–1462.

Tan, S.-I., Liu, Z., Tran, V.G., Martin, T.A., Zhao, H. 2025. *Issatchenkia orientalis* as a platform organism for cost-effective production of organic acids. Metabolic Engineering.

Thorwall, S., Schwartz, C., Chartron, J.W., Wheeldon, I. 2020. Stress-tolerant non-conventional microbes enable next-generation chemical biosynthesis. Nature Chemical Biology, 16(2), 113–121.

Tran, V.G., Mishra, S., Bhagwat, S.S., Shafaei, S., Shen, Y., Allen, J.L., Crosly, B.A., Tan, S.-I., Fatma, Z., Rabinowitz, J.D. 2023. An end-to-end pipeline for succinic acid production at an industrially relevant scale using *Issatchenkia orientalis*. Nature Communications, 14(1), 6152.

Tsugawa, H., Cajka, T., Kind, T., Ma, Y., Higgins, B., Ikeda, K., Kanazawa, M., VanderGheynst, J., Fiehn, O., Arita, M. 2015. MS-DIAL: data-independent MS/MS deconvolution for comprehensive metabolome analysis. Nature Methods, 12(6), 523–526.

Väisänen, T., Kilpeläinen, P., Kitunen, V., Lappalainen, R., Tomppo, L. 2019. Effect of steam treatment on the chemical composition of hemp (*Cannabis sativa* L.) and identification of the extracted carbohydrates and other compounds. Industrial Crops and Products, 131, 224–233.

Vo, T.M., Park, S. 2024. High-yield β-alanine production from glucose and acetate in *Escherichia coli*. Biotechnology and Bioprocess Engineering, 1–11.

Volk, M.J., Tran, V.G., Tan, S.-I., Mishra, S., Fatma, Z., Boob, A., Li, H., Xue, P., Martin, T.A., Zhao, H. 2022. Metabolic engineering: methodologies and applications. Chemical Reviews, 123(9), 5521–5570.

Wang, K., Wu, Z., Du, J., Liu, Y., Zhu, Z., Feng, P., Bi, H., Zhang, Y., Liu, Y., Chen, B. 2023. Metabolic Engineering of *Saccharomyces cerevisiae* for Conversion of Formate and acetate into free fatty acids. Fermentation, 9(11), 984.

Wang, X., Hou, J., Cui, J., Wang, Z., Chen, T. 2024. Engineering *Corynebacterium glutamicum* for the efficient production of 3-hydroxypropionic acid from glucose via the β-alanine pathway. Synthetic and Systems Biotechnology.

Wang, Y., Wang, Y., Cui, J., Wu, C., Yu, B., Wang, L. 2025. Non-conventional yeasts: promising cell factories for organic acid bioproduction. Trends in Biotechnology.

Wu, Z.-Y., Sun, W., Shen, Y., Pratas, J., Suthers, P.F., Hsieh, P.-H., Dwaraknath, S., Rabinowitz, J.D., Maranas, C.D., Shao, Z. 2023. Metabolic engineering of low-pH-tolerant non-model yeast, *Issatchenkia orientalis*, for production of citramalate. Metabolic Engineering Communications, 16, e00220.

Xi, Y., Zhan, T., Xu, H., Chen, J., Bi, C., Fan, F., Zhang, X. 2021. Characterization of JEN family carboxylate transporters from the acid□tolerant yeast *Pichia kudriavzevii* and their applications in succinic acid production. Microbial Biotechnology, 14(3), 1130–1147.

Xiao, H., Shao, Z., Jiang, Y., Dole, S., Zhao, H. 2014. Exploiting *Issatchenkia orientalis* SD108 for succinic acid production. Microbial Cell Factories, 13, 1–11.

Zhang, B., Li, R., Yu, L., Wu, C., Liu, Z., Bai, F., Yu, B., Wang, L. 2023. L-lactic acid production via sustainable neutralizer-free route by engineering acid-tolerant yeast *Pichia kudriavzevii*. Journal of Agricultural and Food Chemistry, 71(29), 11131–11140.

Zhang, Y., Zabed, H.M., Yun, J., Zhang, G., Wang, Y., Qi, X. 2021. Notable improvement of 3-hydroxypropionic acid and 1,3-propanediol coproduction using modular coculture engineering and pathway rebalancing. ACS Sustainable Chemistry & Engineering, 9(12), 4625–4637.

Zhou, S.-P., Ke, X., Jin, L.-Q., Xue, Y.-P., Zheng, Y.-G. 2024. Sustainable management and valorization of biomass wastes using synthetic microbial consortia. Bioresource Technology, 130391.

