## Supplementary material for "Acetate metabolism during xylose fermentation enhances 3-hydroxypropionic acid production in engineered acid-tolerant *Issatchenkia orientalis*": Table S1, Fig. S1, Fig. S2, Fig. S3

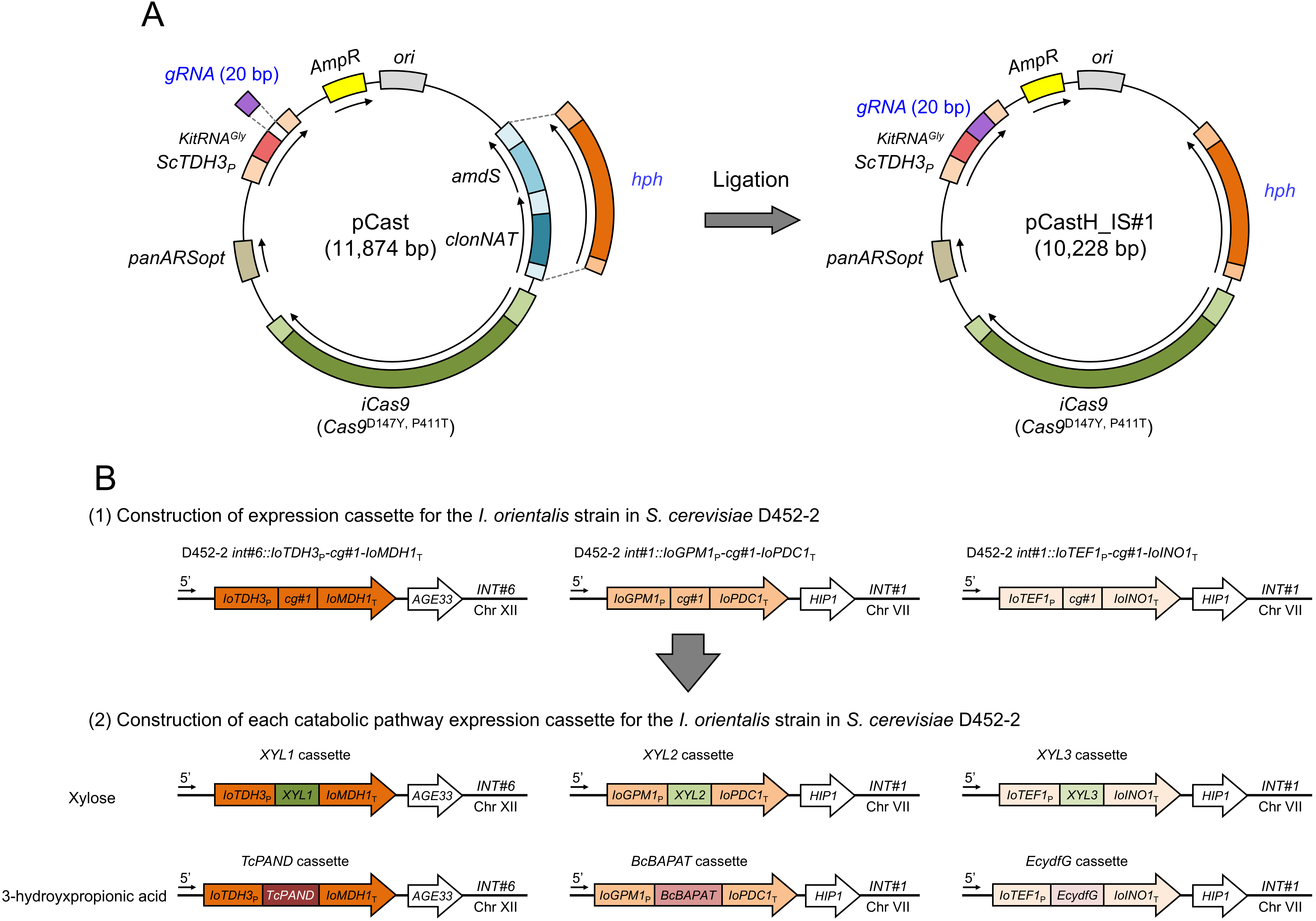

**Fig. S1. Development of a Cas9-based genetic engineering platform for *Issatchenkia orientalis* strain.** (A) Schematic of the construction strategy for the pCastH_IS#1 plasmid containing a hygromycin B resistance marker (*hph*). (B) Construction of expression cassettes for *I. orientalis* using the *Saccharomyces cerevisiae* D452-2 strain.

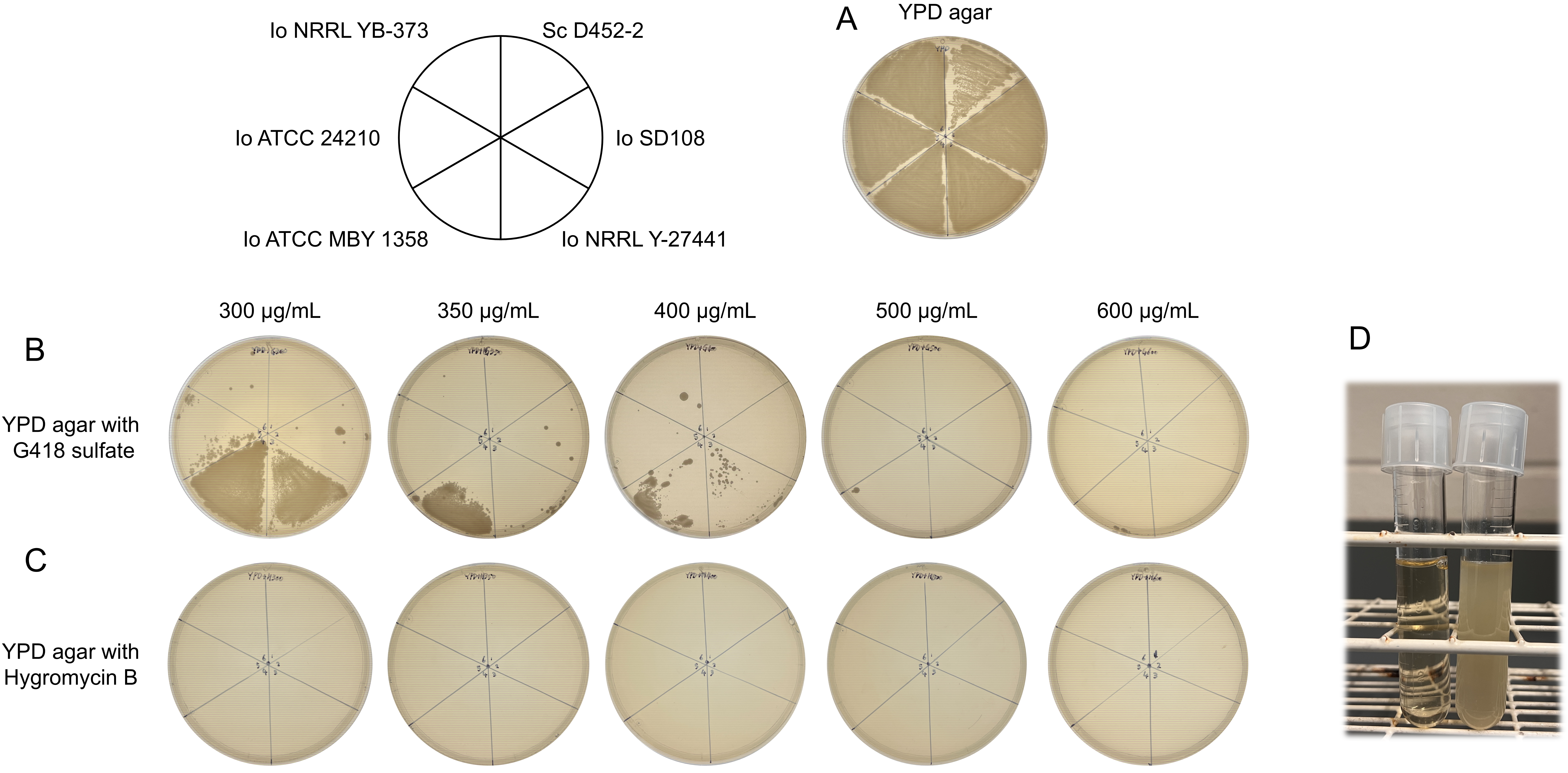

**Fig. S2. Antibiotic resistance profiles of *S. cerevisiae* (Sc) and five *I. orientalis* (Io) strains.** (A-C) Wild-type *S. cerevisiae* D452-2 and five wild-type *I. orientalis* strains (SD108, NRRL Y-27441, ATCC MBY 1358, ATCC 24210, NRRL YB-373) were grown on YPD agar (A), or YPD agar supplemented with increasing concentrations of G418 sulfate (B; 300, 350, 400, 500, 600 μg/mL) or hygromycin B (C; 300, 350, 400, 500, 600 μg/mL). (D) Growth of wild-type *I. orientalis* NRRL Y-27441 strains cultured in YPD broth containing 300 μg/mL hygromycin B at 30 ℃ for 24 h, without (left tube) or with (right tube) the pCastH plasmid.

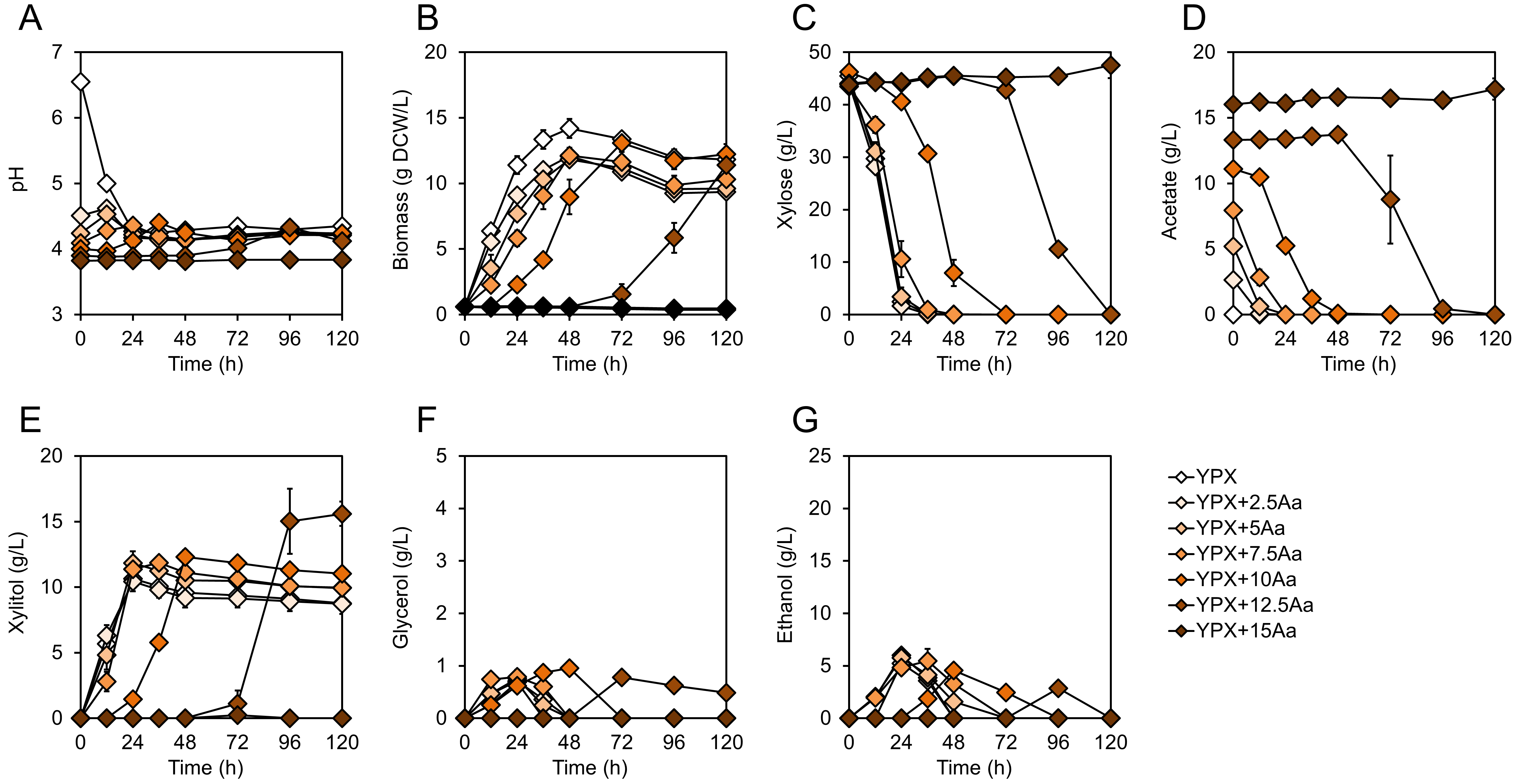

**Fig. S3. Fermentation performance of the IoDY01H strain in xylose and acetate co-utilization.** (A-G) Time-course profiles of pH (A), cell growth (B), substrates consumption—xylose (C) and acetate (D)—and by-product formation, including xylitol (E), glycerol (F), and ethanol (G), during fermentation with 40 g/L D-xylose and varying acetate concentrations (0, 2.5, 5.0, 7.5, 10, 12.5, and 15 g/L). Fermentations were conducted under aerobic conditions (250 rpm) with an initial cell density of 0.6 g DCW/L. Data represent biological triplicates, and error bars indicate standard deviations.

**Table S1.** Primers and guide RNA sequences used in this study.

| **Primers** | **Sequences (5’-)^1)^** | **Description** |
| --- | --- | --- |
| OH554 | CACCTTTCGAGAGGACGATG | Vector_pCast_F |
| OH555 | CCGCACGGTTATCCACAG | Vector_pCast_R |
| OH556 | TTCTGTGGATAACCGTGCGGCAGCGACATGGAGGCCCA | Insert_Hyg_F |
| OH557 | CATCGTCCTCTCGAAAGGTGCGTCCCAAAACCTTCTCAAGC | Insert_Hyg_R |
| OH558 | GGGAAACGCCTGGTATCTTT | Conf_pBR322ori_F |
| OH559 | AAGCAACCATCCTCCGCTC | Conf_AaTEF1P_R |
| OH544 | CGCATAATTTCAACACCTTACTCC | Insert_IS#1_F |
| OH545 | AAACGGAGTAAGGTGTTGAAATTA | Insert_IS#1_R |
| OH737 | CGCAAAGGTTTTGCAACTCCCAAG | Insert_IS#4_F |
| OH738 | AAACCTTGGGAGTTGCAAAACCTT | Insert_IS#4_R |
| OH315 | CGGAGGAGACCGCTATAACCGGTTTGAATTTAATATGGATATGGAGATGAATTTG | Donor_int#6_IoTDH3_P__F |
| OH316 | cctCCGGTGACGGGTAGGTGTACTTTTTGTAATTGTGTTTGTTTGTG | Donor_int#6_IoTDH3_P__R |
| OH317 | GTACACCTACCCGTCACCGGaggAGGTGAAACACAACAACC | Donor_int#6_IoMDH1_T__F |
| OH318 | ATGAACTTGCTTGCTGTCAAACTTCTGAGTTGCGTACAGGGTTATAAAGTTG | Donor_int#6_IoMDH1_T__R |
| OH319 | GAGAAGTTTTTTTACCCCTCTCCACAGATCCGAAAAATGCACCACACC | Donor_int#1_IoGPM1_P__F |
| OH320 | cctCCGGTGACGGGTAGGTGTACTTTGTGTGTGTGTTTTAAGATATAC | Donor_int#1_IoGPM1_P__R |
| OH321 | GTACACCTACCCGTCACCGGaggTGACATCTGAATGTAAAATGAAC | Donor_int#1_IoPDC1_T__F |
| OH322 | CCGGGTAGATTTTTCCGTAACCTTGGTGTCTTGATGGATTGTTTTAGTTTTTA | Donor_int#1_IoPDC1_T__R |
| OH323 | GAGAAGTTTTTTTACCCCTCTCCACAGATCTTTGAAACATCATGAAAACTG | Donor_int#1_IoTEF1_P__F |
| OH324 | cctCCGGTGACGGGTAGGTGTACTGTGATATATAAGTTAGATTTGTC | Donor_int#1_IoTEF1_P__R |
| OH325 | GTACACCTACCCGTCACCGGaggCTACAACAAGATGTTTGTTC | Donor_int#1_IoINO1_T__F |
| OH326 | AGGTAGACCGGGTAGATTTTTCCGTAACCTTGGTGTCATCTGTCAACAACGTACTG | Donor_int#1_IoINO1_T__R |
| OH334 | AAAACACACAAAACACACAAACAAACACAATTACAAAAAATGCCTTCTATTAAGTTGAAC | Donor_IoTDH3_P__XYL1_F |
| OH335 | TTCAAGCTAAAAAAGGAGGTTGTTGTGTTTCACCTTTATTAGACGAAGATAGGAATCTTG | Donor_IoMDH1_T__XYL1_R |
| OH336 | AAGAATCACGTACAATTGTATATCTTAAAACACACACACAAAATGACTGCTAACCCTTCC | Donor_IoGPM1_P__XYL2_F |
| OH337 | GTAATTCATTTTAATGTTCATTTTACATTCAGATGTCATTATTACTCAGGGCCGTCAATG | Donor_IoPDC1_T__XYL2_R |
| OH338 | TTTTCCTTCAACAGACAAATCTAACTTATATATCACAAAAAATGACCACTACCCCATTTG | Donor_IoTEF1_P__XYL3_F |
| OH339 | ACAAGTTGCTCCCCTTGAACAAACATCTTGTTGTAGTTATTAGTGTTTCAATTCACTTTC | Donor_IoINO1_T__XYL3_R |
| OH783 | AAAACACACAAAACACACAAACAAACACAATTACAAAAAAAAAATGCCTGCTACTGGTG | Donor_IoTDH3_P__TcPAND_F |
| OH784 | CAAGCTAAAAAAGGAGGTTGTTGTGTTTCACCTTTATTACAAATCAGATCCTAATCTTTC | Donor_IoMDH1_T__TcPAND_R |
| OH785 | CGTACAATTGTATATCTTAAAACACACACACAAAAAAAATGGAATTGATGATTGTTCAAG | Donor_IoGPM1_P__BcBAPAT_F |
| OH786 | AATTCATTTTAATGTTCATTTTACATTCAGATGTCATTATTACAATTGAGCCAAACATTC | Donor_IoPDC1_T__BcBAPAT_R |
| OH787 | TCCTTCAACAGACAAATCTAACTTATATATCACAAAAAATGATCGTTTTAGTAACTGGAG | Donor_IoTEF1_P__EcydfG_F |
| OH411 | CTATCGGCCTCTTTTTCTCCGGGTGTGGTGCATTTTTCGCGTACAGGGTTATAAAGTTG | Donor_IoMDH1_T_-IoGPM1_P__R |
| OH412 | CTACTTTAGATGCTCCTCTGAACAACTTTATAACCCTGTACGCGAAAAATGCACCACACC | Donor_IoGPM1_P_-IoMDH1_T__F |
| OH413 | CACAGAGGGTGAAACAGTTTTCATGATGTTTCAAATTGATGGATTGTTTTAGTTTTTA | Donor_IoPDC1_T_-IoGPM1_P__R |
| OH414 | ATATATAATTTTATAATAAAAACTAAAACAATCCATCAATTTGAAACATCATGAAAACTG | Donor_IoTEF1_P_-IoPDC1_T__F |
| OH619 | ATGAACTTGCTTGCTGTCAAACTTCTGAGTTGATCTGTCAACAACGTACTG | Donor_int#6_IoINO1_T__R |
| OH410 | AAAATATTGAACCGTCGAAACGTCCCAAACAAGGAAAATATGGATATGGAGATGAATTTG | Donor_is#1_IoTDH3_P__F |
| OH415 | AAGGTGAGAATTCAAAATGTTATTTTTGATCATGTAAAGGGATCTGTCAACAACGTACTG | Donor_is#1_IoINO1_T__R |
| OH774 | CATGCTTTACAAGATGATAAATCTATTTATTGCTCATATGGATATGGAGATGAATTTG | Donor_is#4_IoTDH3_P__F |
| OH744 | ATTTGCTTCTCACTATTGCTTGCTTGTACTCAAAGAGCAAAATCTGTCAACAACGTACTG | Donor_is#4_IoINO1_T__R |
| OH196 | GGTTCTGACTCCTACTGAGC | Conf_INT#6_F |
| OH197 | AGCATCGAGTACGGCAGTTC | Conf_INT#6_R |
| OH276 | GTGAGTTCTCATAACCTCG | Conf_INT#1_F |
| OH277 | CTAGTGCGCCAAGAGAGTAATT | Conf_INT#1_R |
| OH427 | TCCTCCTCCGCCACTAAAGTC | Conf_IoTDH3_P__R |
| OH331 | AAGCGAGGAAATGAGCGAC | Conf_IoTDH3_P__F |
| OH416 | AAAGCCCAAGAAGGAGAAACC | Conf_IoGPM1_P__R |
| OH417 | TTTCGTTTCTCCCTTCCCTGATAG | Conf_IoPDC1_T__F |
| OH418 | GTGCAGTTGAGAGGTGGGAG | Conf_IoTEF1_P__R |
| OH419 | GTCTATCTCTCCCTATCGCTCTG | Conf_IoINO1_T__F |
| OH242 | GAGCATAGGCATAGAGAAGCCGTC | Conf_IS#1_F |
| OH247 | ACTAATTCCAGTTGAGATACTCACGCACAAGA | Conf_IS#1_R |
| OH745 | ATGAGTTCAGCCGATACTAGAGC | Conf_IS#4_F |
| OH746 | GGTGTGGCCTTCAAATGATCC | Conf_IS#4_R |
| **gRNA** | **guide RNA and PAM sequences (5’-)** | **References** |
| For *Saccharomyces cerevisiae* | | |
| *cg#1* | GTACACCTACCCGTCACCGG AGG | (Jeong et al., 2024) |
| *INT#1* | GATACTTATCATTAAGAAAA TGG | (Jeong et al., 2020a) |
| *INT#6* | TTGTCACAGTGTCACATCAG CGG | (Jeong et al., 2020a) |
| For *Issatchenkia orientalis* | | |
| *IS#1* | TAATTTCAACACCTTACTCC TGG | (Lee et al., 2022b) |
| *IS#4* | AAGGTTTTGCAACTCCCAAG CGG | This study |

^1)^ The flanking region is underlined.
